# NOX4 links metabolic regulation in pancreatic cancer to endoplasmic reticulum redox vulnerability and dependence on PRDX4

**DOI:** 10.1101/2020.10.08.330035

**Authors:** Pallavi Jain, Anna Dvorkin-Gheva, Erik Mollen, Michael Xie, Fatima Jessa, Piriththiv Dhavarasa, Stephen Chung, Kevin R. Brown, Gun Ho Jang, Faiyaz Notta, Jason Moffat, David Hedley, Paul C. Boutros, Bradly G. Wouters, Marianne Koritzinsky

## Abstract

Pancreatic ductal adenocarcinoma (PDAC) is an almost universally fatal malignancy and there is an urgent need for new therapeutic targets. PDAC cells harbor genetic alterations and display metabolic changes that render them vulnerable to perturbations in redox homeostasis. Peroxiredoxin 4 (PRDX4) supports redox homeostasis by metabolizing H_2_O_2_ in the endoplasmic reticulum (ER). Based on functional genomics, we found that PDAC cell lines are dependent on PRDX4 for their growth and survival. We validated this dependency in established and primary PDAC cells, as well as 3D models and orthotopic xenografts. Cell death induced by PRDX4 depletion was accompanied by increased levels of reactive oxygen species (ROS), DNA damage and a DNA-PKcs-governed DNA damage response. As such, PRDX4 depletion also sensitized cells and tumors to ionizing radiation. The source of ROS that created a dependency on PRDX4 was attributed to NADPH oxidase 4 (NOX4), which localizes to the ER membrane. The functional requirement for PRDX4 was correlated with cellular NADPH levels across different models of PDAC and could be rescued by depletion of NOX4 or NADPH. As such, this study has identified NOX4 as a link between metabolic deregulation and ER-specific redox vulnerability in PDAC. Since PRDX4 is not an essential gene in normal tissues, our work also suggests that PRDX4 represents a novel therapeutic target for pancreatic cancer that may be particularly potent in combination with radiotherapy.

## Introduction

Pancreatic ductal adenocarcinoma (PDAC) accounts for more than 80% of pancreatic cancer cases and is associated with an overall 5-year survival rate of only 8%. Although the efficacy of surgery, chemo- and radiation therapies have improved over the last decade, immunotherapy response has been disappointing and other targeted personalized therapies are lacking (1).

PDAC has a complex landscape of genetic alterations with prevalent chromothripsis and mutations in *KRAS*, *TP53*, *SMAD4* and *CDKN2A* (2–4). From the products of these lesions, only the specific KRAS^G12C^ mutation is potentially actionable at this point, with a covalent inhibitor currently in clinical trials (5). However, heterogeneous and complex compositions of genetic alterations may in concert drive common phenotypes that expose specific vulnerabilities. One such phenotype that has emerged as a potential vulnerability in cancer is aberrant redox homeostasis (6). The high proliferation rates of cancer cells are accompanied by production of reactive oxygen species, which can become cytotoxic. Cancer cells have adapted to tolerate the exacerbated oxidative burden by increasing the cellular antioxidant capacity through transcriptional upregulation of antioxidant enzymes and metabolic reprogramming (6).

In pancreatic cancer, systems approaches comprising transcriptomic and metabolomic analysis have revealed that *KRAS*^G12D^ stimulates glucose uptake, hexosamine biosynthesis and diversion of glycolysis intermediates to the non-oxidative arm of the pentose phosphate pathway (PPP) for ribose biosynthesis (7). Bypass of the oxidative arm of the PPP is perhaps puzzling, given its important role in providing NADPH important for macromolecule biosynthesis and ROS detoxification. However, several *KRAS*-driven mechanisms contribute to support redox homeostasis in other ways. *KRAS*^G12D^ promotes transcription of *NFE2L2*, which is a master transcriptional regulator of antioxidant enzymes (8). Oncogenic *KRAS* is also responsible for diverting mitochondrial glutamine-derived metabolites to cytosolic malate production which through malic enzyme 1 (ME1) is converted to pyruvate in an NADPH-producing reaction (9). This reprogramming was shown to be important for maintaining redox homeostasis and viability of PDAC cells. As such, glutamine metabolism emerged as a potential therapeutic target for pancreatic cancer (10).

Another potential therapeutic strategy could be to target redox balance in more direct ways. *In vivo* studies on development of PDAC show that KRAS-dependent ROS are induced in acinar cells, gradually increase during acinar-to-ductal metaplasia and pancreatic intraepithelial neoplasia (PanIN) formation, and are essential for tumorgenicity (11–13). However, although a certain level of oxidative stress promotes cancer progression, excessive ROS can also modify and damage macromolecules including lipids, proteins and DNA, and thereby evoke toxicity (14). ROS can be generated in multiple organelles including the mitochondria, ER, and peroxisomes by enzymatic reactions involving cyclooxygenases, oxidoreductases, NADPH oxidases (NOXs), xanthine oxidases and lipoxygenases and through the iron-catalyzed Fenton reaction (6). To equilibrate this, ROS scavenging defense systems are present in the form of superoxide dismutases (SODs), glutathione peroxidase (GPX), peroxiredoxins, thioredoxin and catalase (6). Several of these systems have been targeted for the benefit of pancreatic and other cancer treatment, although with limited success, likely because of limited therapeutic index caused by normal tissue toxicity (15). Peroxiredoxins (PRDXs) are a ubiquitous and potentially druggable family of antioxidant proteins that have heterogeneous tissue and intracellular distributions, which may confer less toxicity upon therapeutic targeting. PRDXs exist as (do-)decamers consisting of dimeric units which metabolize hydrogen peroxide (H_2_O_2_) (16, 17). There are six PRDX family members with PRDX1 and PRDX6 localized to the cytosol, while PRDX2, PRDX3, PRDX4 and PRDX5 are localized to the nucleus, mitochondria, endoplasmic reticulum and peroxisomes, respectively (18, 19).

Peroxiredoxins contain an exposed peroxidatic cysteine, which in the catalytic cycle is oxidized to sulfenic acid by H_2_O_2_. PRDX1-5 contain another resolving cysteine (C_R)_ that subsequently reacts with CP to form an intra- or inter-molecular disulfide bond, preventing further irreversible hyperoxidation. The disulfide bond is a substrate for reduction by thioredoxin or thioredoxin-like proteins, resulting in recycling of PRDXs back to their reactive thiol state. Through this mechanism, PRDX family members are thought to scavenge more than 90% of cellular peroxides (17).

To explore the essentiality of PRDX family members in pancreatic cancer, we exploited a functional genomics dataset. We demonstrate that approximately half of primary and established pancreatic cell lines are dependent on PRDX4 for their proliferation and survival *in vitro* and *in vivo*. We also demonstrate that this dependency is driven by cellular compartment-specific metabolism and is associated with severe levels of DNA damage, which can be exploited therapeutically.

## Results

### Essentiality of peroxiredoxins

To determine the essentiality of PRDX family members in pancreatic cancer cell lines, we mined data from published functional genomics screens that used a lentiviral-based genome-wide pooled RNA interference library targeting 16,056 protein coding genes with 78,432 short hairpin (sh) RNAs (20) (Supplementary Figure 1A). All 36 cell lines were sensitive to losing at least one PRDX family member, while a vast majority of cell lines were sensitive to the depletion of several PRDXs (Figure 1A). PRDX3 was the most frequently essential protein in the family with 85% of cell lines sensitive to its depletion. There was no co-dependence in sensitivity to depletion between any pairs of PRDX family members (Fisher exact tests, all p-values n.s.). Taken together, these observations suggest that essentiality is associated with specific organelle vulnerabilities in each cell line, rather than a general sensitivity to loss of anti-oxidant capacity. RNA sequencing (RNAseq) from 32 cell lines revealed that dependency on individual PRDX family members was not associated with its own mRNA abundance across cell lines (Supplementary Figure 1B), indicating that other genetic or epigenetic features confer PRDX addiction.

**Figure 1:**
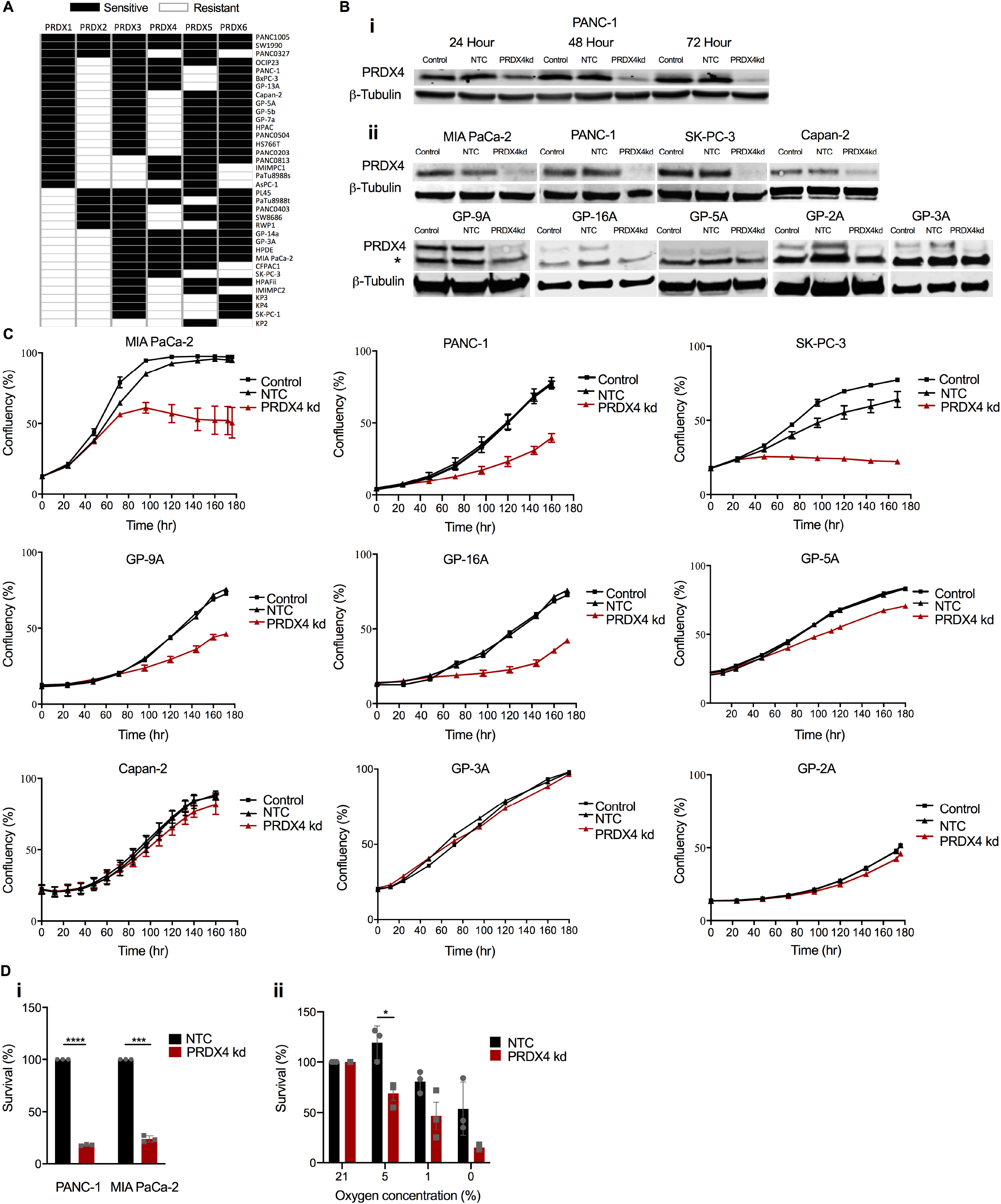
Pancreatic cancer cell lines depend on PRDX4 for growth and survival. A) Illustration of cell lines (n=36) where a PRDX was deemed essential (black boxes) in a functional genomics screen (20). B) i) Western blot showing PRDX4 expression at indicated time points after siRNA transfection in PANC-1 cells. ii) Western blot showing PRDX4 expression in established (PANC-1, MIA PaCa-2, SK-PC-3, Capan-2) cell lines and patient derived primary cell lines (GP-9A, GP-16A, GP-5A, GP-3A and GP-2A) 72 hours after transfection with siRNA targeting nothing (no-template control, NTC) or PRDX4 (PRDX4kd). * denotes a cross-reacting band. C) Cell confluency as a function of time after siRNA transfection as measured by automated live cell imaging for the indicated cell lines. Data points represent the average of 3 biological replicates ± SD. D) Percent survival determined by clonogenic assay for MIA PaCa-2 and PANC-1 cells. i) Cells were plated 72 hours after siRNA transfection and survival was normalized to the plating efficiency of NTC. ii) Cells were plated 72 hours after siRNA transfection with exposure to the indicated percent oxygen for the last 24 hours, and survival was normalized to the plating efficiency of each group in air. Data points represent independent experiments and bars the average ± SEM. ***: p<0.005; **: p<0.01; * p<0.05.

The results outlined above suggest that depletion of PRDX family members may represent a vulnerability in pancreatic cancer cells. However, the ubiquitous expression of most *PRDX*s and the severe phenotypes of *Prdx1, Prdx2, Prdx3*, *Prdx5 and Prdx6* knockout mice raise concerns that targeting these family members may be associated with toxicities (21–25). Therefore, we focused on *PRDX4,* which is mainly expressed in the liver and pancreas, is not associated with severe phenotypes when knocked out in a mouse model, but was required for the proliferation of 47% of pancreatic cancer cell lines (Figure 1A) (26).

PRDX4 is localized in the ER where it protects cells against oxidative stress (27, 28). Analyzing RNAseq data from 195 advanced pancreatic cancer patients collected as part of the multi-institutional Canadian COMPASS trial (Comprehensive Molecular Characterization of Advanced Pancreatic Ductal Adenocarcinomas (PDAC) for Better Treatment Selection: A Prospective Study, NCT02750657), we found that high *PRDX4* expression was associated with the more aggressive basal-like subtype, compared to the less aggressive classical subtype (p=0.024)(Supplementary Figure 2A) (29, 30). Expression of PRDX4 mRNA was also higher in biopsies collected from liver metastases than those collected from localized tumors in the pancreas (p=1.957e-9)(Supplementary Figure 2B). Moreover, high *PRDX4* expression was correlated with shorter overall survival in a cohort of 178 patients with resectable tumors (p=0.028) (Supplementary Figure 2C). Together these data demonstrate that high PRDX4 mRNA abundance is associated with more aggressive disease.

### PRDX4 is essential in established and primary pancreatic cancer cell lines

To validate the results of the shRNA screening dataset, we tested the ability of independent siRNA sequences targeting PRDX4 to inhibit the proliferation and survival of pancreatic cancer cells. Efficient knockdown of PRDX4 could be achieved 72 hours after transfection of PANC-1 cells (Figure 1B i), in line with the reported long half-life of PRDX4 protein (31). Efficient knockdown was also validated in three additional pancreatic cancer cell lines (MIA PaCa-2, SK-PC-3, Capan-2) (Figure 1B ii) as well as in five primary pancreatic cancer cell lines (GP-2, GP-3A, GP-5A, GP-9A, GP-16A) (Figure 1B ii). Depleting PRDX4 significantly reduced proliferation of MIA PaCa-2, PANC-1 and SK-PC-3 cells, whereas Capan-2 cells did not show any change in proliferation (Figure 1C), confirming the results obtained in the functional genomics dataset. Importantly, PRDX4 depletion also significantly repressed the proliferation of three primary pancreatic cell lines (GP-9A, GP-16A, GP-5A), while two were resistant (GP-3A, GP-2A) (Figure 1C), recapitulating the frequency of essentiality in established cell lines. To assess the long-term impact of transient PRDX4 depletion, we plated cells for colony formation 96 hours after siRNA transfection. Transient PRDX4 depletion reduced colony formation by ~80% in both cell lines assessed (p<0.001)(Figure 1D i). The tumor microenvironment is characterized by oxygen gradients and hypoxia, which drives aggressive tumor phenotypes, treatment resistance and poor patient outcomes (32). To assess whether oxygen availability affects PRDX4 dependency, we exposed cells to various oxygen concentrations during PRDX4 depletion, and plated cells for colony formation. Depletion of PRDX4 was more toxic to cells under hypoxic conditions (Figure 1D ii). These results demonstrate that *PRDX4* is a required gene for proliferation and survival in a subset of established and primary PDAC cell lines, a dependency that increases in reduced oxygen environments.

### PRDX4 dependency in 3D and *in vivo* tumor models

To determine the long-term effects of continuous PRDX4 depletion, we generated two PDAC cell lines (MIA PaCa-2 and PANC-1) expressing doxycycline (Dox) inducible shRNA targeting PRDX4 (shPRDX4). Dox exposure reduced PRDX4 abundance (Figure 2A) and inhibited proliferation (Figure 2B) in both cell lines. Since cells cultured in 2D can undergo cytoskeletal rearrangements and acquire artificial polarity (33), we also assessed the effect of loss of PRDX4 in a 3D spheroid model. Treatment with Dox to deplete PRDX4 for 10 days significantly inhibited growth and resulted in spheroid shrinkage (Figure 2C).

**Figure 2:**
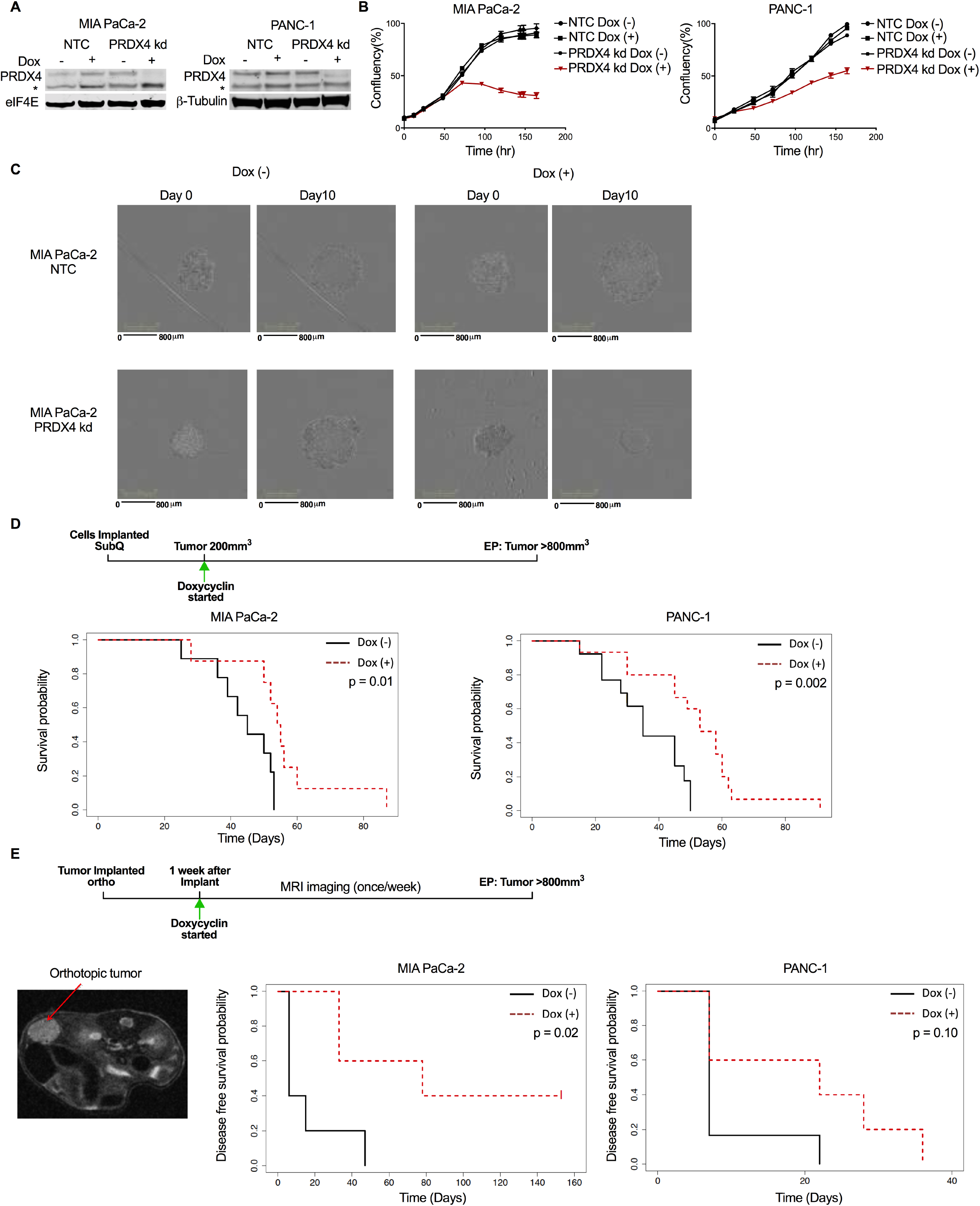
PRDX4 depletion inhibits proliferation of spheroids and xenografts. A) Western blots showing PRDX4 expression in MIA PaCa-2 and PANC-1 cells expressing doxycycline (Dox)-inducible shRNA targeting nothing (NTC) or PRDX4 (PRDX4 kd), following treatment with Dox for 72 hours. * denotes a cross-reacting band. B) Cell confluency as a function of time for the same cells after addition of Dox as measured by automated live cell imaging for the indicated cell lines. Data points represent the average of 3 biological replicates ± SD. C) Micrographs of spheroids made from MIA PaCa-2 cells at the start of Dox exposure and 10 days thereafter. D) Survival probability of mice carrying subcutaneous MIA PaCa-2 (n=17) or PANC-1 (n=27) tumors, where the cells expressed Dox-inducible shRNA targeting PRDX4. Mice were fed Dox in chow from the day tumors reached 200mm^3^ (Day 0) until endpoint (EP) of >800mm^3^ tumor burden. E) Mice were implanted orthotopically with MIA PaCa-2 (n=11) or PANC-1 (n=11) cells expressing Dox-inducible shRNA targeting PRDX4 and received Dox in chow from 1 week thereafter (Day 0). Tumors were monitored with MR imaging, an example of which is shown to the left, until endpoint (EP) of >800mm^3^ tumor burden. Right: disease free survival probability of mice as a function of time after Dox exposure.

To assess the effect of PRDX4 depletion on tumor growth *in vivo*, we established subcutaneous xenografts from MIA PaCa-2 and PANC-1 cells expressing Dox-inducible shPRDX4 in immune-compromised mice (NOD-Rag1^null^/IL2rg^null^). To mimic a therapeutic situation, we started treating mice with Dox after tumors were established and had reached a size of 200mm^3^ (Figure 2D). We measured expression of *PRDX4* when mice reached endpoint, as well as in a separate cohort of mice after one week of Dox treatment. One week of Dox significantly reduced the abundance of PRDX4 mRNA in both tumor models (p<0.01), with undetectable levels in many tumors (Supplementary Figure 3A). At endpoint, PRDX4 mRNA was detectable but still low in all MIA PaCa-2 tumors, while knockdown appeared less efficient in PANC-1 tumors (Supplementary Figure 3A). This suggests that Dox treatment conferred selection of cells resistant to shPRDX4 through transgene silencing or other mechanisms. Nevertheless, targeting PRDX4 increased the survival of mice bearing tumors from both cell models (p=0.01 and p=0.002, respectively) (Figure 2D and Supplementary figure 2B).

Tumors grown at the orthotopic site tend to better recapitulate the patient tumor micro-environment and provide more exposure to relevant metastatic routes (34). We therefore implanted the same models in the mouse pancreas and started Dox treatment 1 week thereafter. We monitored tumor growth with MRI imaging (Figure 2E). Mice presented with tumors in the pancreas as well as a heterogenous distribution of metastases to distant sites such as lymph nodes, spleen and liver. Treatment with Dox significantly increased the disease-free survival of mice implanted with MIA PaCa-2 tumors (p=0.02), with 40% of the mice remaining tumor-free (Figure 2E and Supplementary figure 2C). A trend towards increased disease-free survival was also observed in mice carrying PANC-1 tumors, but this was not statistically significant, possibly due to outgrowth of shRNA-resistant clones. Taken together, these findings demonstrate that PRDX4 targeting can inhibit tumor growth *in vitro* and *in vivo*, but that similar to other targeted agents, adaptation and selection mechanisms may affect long-term response.

### PRDX4 dependency is associated with increased ROS

We next sought to identify the mechanism underlying PRDX4 dependency. PRDX4 is localized to the ER where its canonical role is to metabolize H_2_O_2_ and thereby protect the ER lumen from oxidative damage (28, 35). We therefore investigated whether there was an association between PRDX4 dependency and ROS levels across two cell lines sensitive (MIA PaCa-2 and PANC-1) and two cell lines resistant (Capan-2 and GP-3A) to PRDX4 depletion. Interestingly, PRDX4 depletion only increased the abundance of ROS in the sensitive cell lines (Figure 3A-B). This indicates that PRDX4 is only required to maintain redox homeostasis in a subset of cell lines, and that these cells consequentially rely on PRDX4 for proliferation and survival.

**Figure 3:**
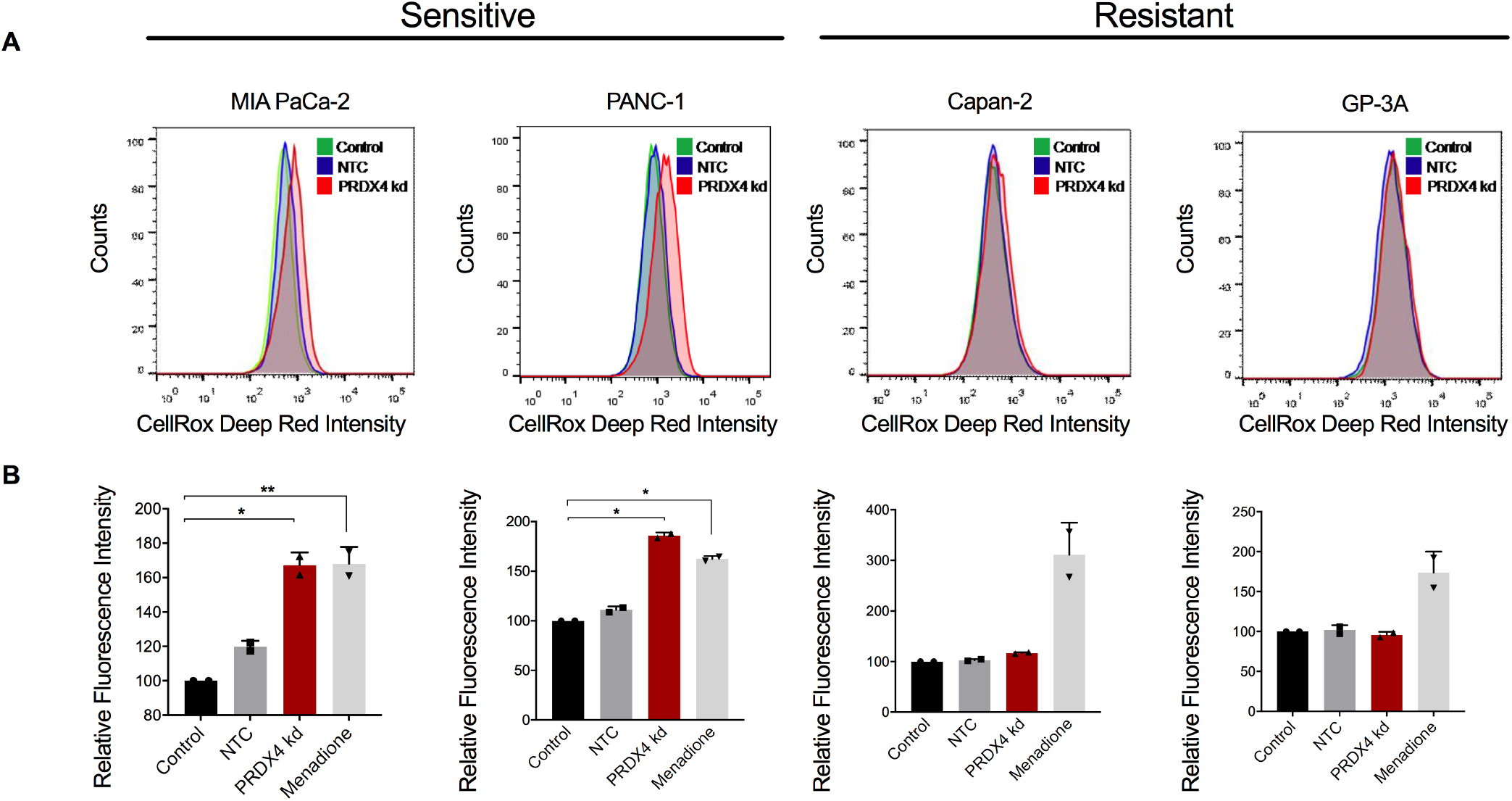
Sensitivity to PRDX4 depletion is associated with increased ROS abundance. A) Flow cytometry histograms of cellular ROS measured by oxidized CellROX in cells sensitive (MIA PaCa-2 and PANC-1) and resistant (Capan-2 and GP-3A) to PRDX4 depletion, 72 hours after transfection with siRNA targeting nothing (NTC) or PRDX4 (PRDX4 kd). B) Quantification of relative fluorescence intensity from samples as shown in A. 100 μM menadione exposure for 1 hour was used as a positive control. Each data point represents an independent experiment; bars represent the average value ± SD. (**: p<0.01; *: p<0.05)

Alterations in the redox environment of the ER can affect protein folding leading to activation of the unfolded protein response (UPR) (36). We therefore examined the expression of UPR-induced transcripts, including XBP1s, ERdj4 and CHOP. There was no detectable change in expression of any of the UPR induced genes upon PRDX4 depletion (Supplementary Figure 4). These observations suggest that the elevated ROS does not cause significant proteotoxicity, and other consequences of loss of redox homeostasis are likely to underlie the dependence on PRDX4.

### Loss of PRDX4 induces DNA damage

Since the ER is in close proximity to the nucleus with continuity between the ER lumen and nuclear intermembrane space, we investigated whether PRDX4 depletion was associated with markers of DNA damage. Indeed, PRDX4 depletion with either siRNA (Figure 4A) or Dox-inducible shRNA (Figure 4B) resulted in phosphorylation of histone H2AX at Ser139 (γH2AX), consistent with the induction of DNA double strand breaks (DSBs). This induction was observed in five established and primary PDAC cell lines where PRDX4 had been deemed essential, but not in three cell lines resistant to PRDX4 depletion (Figure 4A and Supplementary Figure 5A).

**Figure 4:**
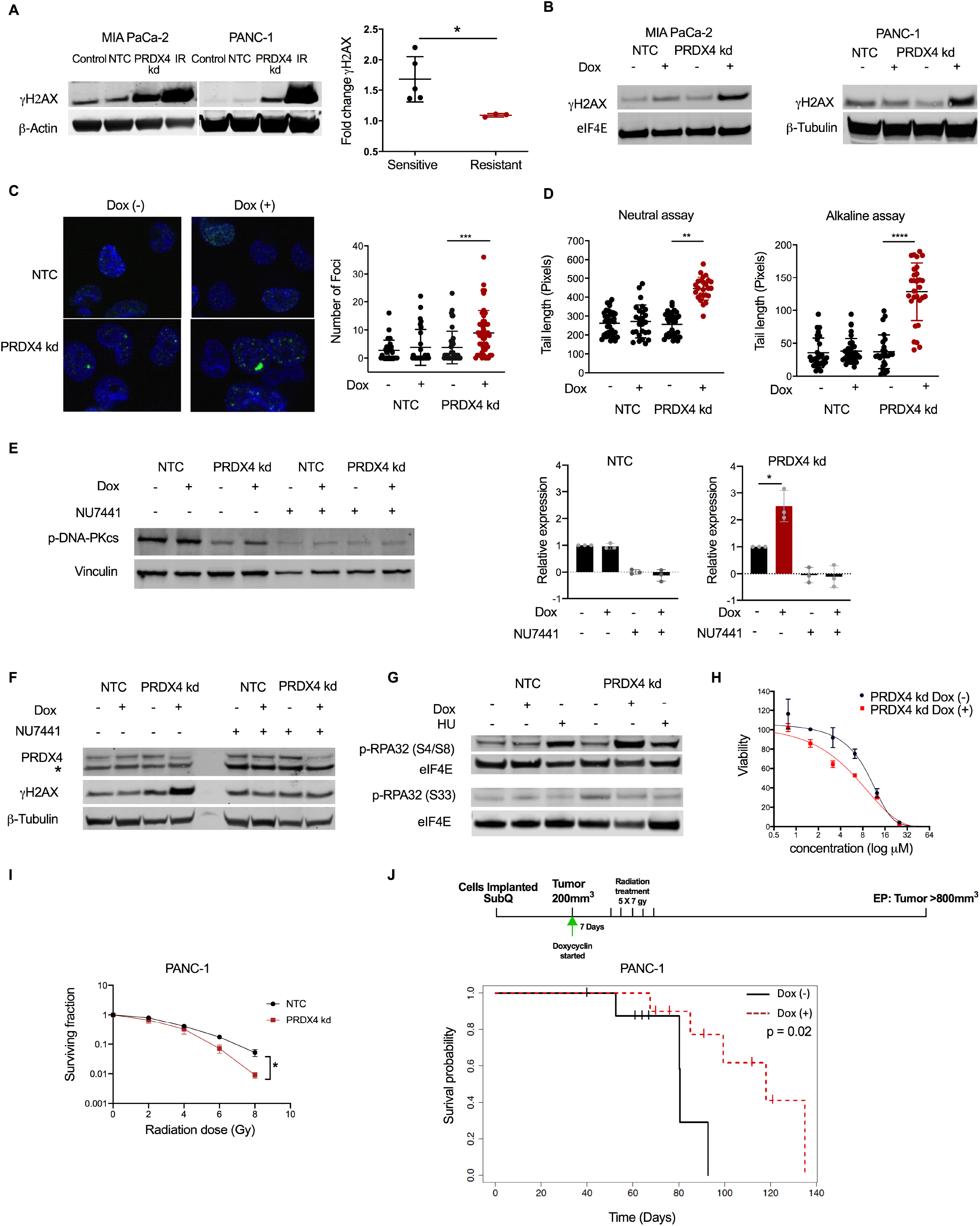
PRDX4 depletion results in a DNA-PK-dependent DNA damage response. A) Western blot showing expression of γH2AX 72 hours after transfection of the indicated cells with siRNA against nothing (NTC) or PRDX4 (PRDX4 kd), or 30 minutes after exposure to 15Gy ionizing radiation (IR). Quantification of fold change in γH2AX expression following transfection with siRNA targeting PRDX4, in cells sensitive (MIA PaCa-2, PANC-1, SK-PC-3, GP-9A and GP-16A) or resistant (Capan-2, GP-2A and GP-3A) to PRDX4 depletion. Each data point represents an individual cell line, the average ± SD is shown. *: p<0.05. B) Western blot showing expression of γH2AX in the indicated cells expressing Dox-inducible shRNA targeting nothing (NTC) or PRDX4 (PRDX4 kd) 72 hours after Dox exposure. C) Fluorescence micrograph of nuclei (DAPI; blue) with γH2AX foci (green) in MIA PaCa-2 cells. Quantification of number of foci per cell. D) Quantification of comet tail length during neutral and alkaline comet assay in MIA PaCa-2 cells. Each data point represents one nucleus, average value is indicated ± SD. ****: p<0.001; ***: p<0.005; **: p<0.01. E) Western blot showing expression of p-DNA-PKcs in MIA PaCa-2 cells after treatment with Dox and/or 10μM NU7441 for 96 hours. Quantification of p-DNA-PKcs expression normalized to the no treatment sample. Each datapoint represents an independent experiment, bars represent the average value ± SD. *: p<0.05. F) Western blot showing expression of γH2AX in MIA PaCa-2 cells after treatment with Dox and/or 10μM NU7441 for 96 hours. * denotes a cross reacting band. G) Western blot showing expression of p-RPA32 (S4/S8) and p-RPA32 (S33) in MIA PaCa-2 cells after treatment with Dox for 96 hours or 5μM hydroxyurea (HU) for 24 hours. H) Viability of cells as in (B) as measured by AlamarBlue reagent after exposure to NU7441 for 24 hours in MIA PaCa-2 cells after treatment with Dox for 96 hours. I) Surviving fraction measured by clonogenic assay of PANC-1 cells plated 72 hours after transfection with siRNA targeting nothing (NTC) or PRDX4 (PRDX4 kd), and 30 minutes after irradiation with the indicated doses. Data points represent the average value from 3 independent experiments ± SEM. J) Survival probability of mice (n=19) carrying subcutaneous tumors from PANC-1 cells expressing Dox-inducible shRNA targeting PRDX4. Dox was administered in chow from the time tumors reached 200mm^3^ (Day 0). Irradiation started 7 days thereafter, with 5 fractions of 7Gy given over 10 days.

H2AX is phosphorylated by phosphoinositide 3-kinase-related kinase (PIKK) family members in megabase regions extending from DSBs, generating a focal γH2AX signal important for regulation of DNA repair (37). We confirmed by immunofluorescence that PRDX4 depletion resulted in an increase in the number of γH2AX foci per cell (p<0.01) (Figure 4C). To confirm that γH2AX foci were a result of the DNA damage response, we quantified DNA damage directly using Comet assays (38). PRDX4 depletion resulted in significantly longer Comet tails when assayed under both alkaline and neutral conditions (Figure 4D), demonstrating the induction of DNA single strand breaks (SSBs) and DSBs, respectively. Consistent with the presence of DNA damage, PRDX4 knockdown resulted in a higher proportion of cells in G2/M phase of the cell cycle, but only in cells lines in which PRDX4 was essential (Supplementary Figure 5B). These data show that PRDX4 depletion results in SSBs, DSBs and a DNA damage response (DDR) in cell lines sensitive to its depletion and suggest that γH2AX could represent a functional biomarker of PRDX4 dependency.

### PRDX4-mediated DDR is driven by DNA-PK

H2AX is a substrate of the PIKK family members ATM, ATR and DNA-PK. To identify the kinase(s) responsible for H2AX phosphorylation upon loss of PRDX4, we examined their activation as reflected by auto-phosphorylation and phosphorylation of their other substrates. Despite the presence of SSBs and DSBs (Figure 4D), we did not detect any signs of activation of ATM or ATR (Supplementary Figure 5C-D). Importantly however, cells were competent in activating ATM and ATR as evidenced by auto- and substrate- (CHK1) phosphorylation after ionizing radiation or hydroxyurea treatment (Supplementary Figure 5C-D). In contrast, we consistently observed a 2-fold increase in phosphorylation of the catalytic subunit of DNA-PK (DNA-PKcs) after PRDX4 depletion (Figure 4E and Supplementary Figure 5E). DNA-PKcs phosphorylation was eliminated in the presence of the selective DNA-PK inhibitor NU7441, demonstrating that it reflects autophosphorylation as expected at this residue (Figure 4E). NU7441 also eliminated γH2AX following PRDX4 knockdown (Figure 4F), consistent with data described above demonstrating that DNA-PK is the only PIKK activated in these models. Furthermore, PRDX4 depletion led to phosphorylation of RPA32 at the DNA-PK-dependent S4/S8 residue, and not at the ATM/ATR-dependent S33 residue (Figure 4G) (39). Although the crosstalk between PIKKs are not fully understood and most certainly are context-dependent, these results suggest that PRDX4 depletion leads to recruitment of primarily DNA-PK to stalled replication forks, resulting in RPA32 and H2AX phosphorylation to mediate repair (40).

We next hypothesized that DNA-PK activation during PRDX4 depletion serves an adaptive role. Consistent with this, we found that PRDX4 knockdown resulted in moderately increased sensitivity to NU7441 (Figure 4H). Further, we speculated that the presence of DNA damage following PRDX4 knockdown could increase sensitivity to ionizing radiation. In line with this, we observed that PANC-1 cells depleted of PRDX4 were sensitized to radiation as measured by their colony-forming ability (Figure 4I). This prompted us to investigate whether combining radiotherapy with PRDX4 depletion *in vivo* could counteract the adaptation/resistance mechanisms that we had observed in previous experiments (Figure 2D-E and Supplementary Figure 3A). Hypo-fractionated radiotherapy schedules are increasingly being used to treat pancreatic cancer in the clinic, so we treated 200mm^3^ PANC-1 xenografts with a similar schedule of 35Gy given in 7Gy fractions over 10 days in the presence or absence of Dox-induced PRDX4 depletion (Figure 4J and Supplementary Figure 5F). PRDX4 targeting significantly improved the outcomes of mice treated with a clinically relevant schedule of radiotherapy (p=0.02).

### PRDX4 dependency requires NOX4

To understand the mechanism underlying PRDX4 dependency, we sought to identify the source responsible for increased ROS upon PRDX4 depletion (Figure 3). One major source of ROS in the ER is endoplasmic oxidoreductin-1-like protein α (ERO1-Lα), which generates H_2_O_2_ when transferring electrons to molecular oxygen from protein disulfide isomerase (PDI) after disulfide bonds are introduced in ER client proteins (41). To determine if the increase in ROS observed upon PRDX4 knockdown in sensitive cells was due to ERO1-Lα, we tested if ERO1-Lα knockdown would rescue the anti-proliferative effects of PRDX4 depletion. However, ERO1-Lα-targeting siRNA offered no protection (Supplementary Figure 6A).

Another potential contributor to ROS production is the trans-membrane NOX4 enzyme complex that produces ROS in a NADPH-dependent manner (42). NOX4 has been reported in several subcellular locations including in the ER membrane where it produces superoxide and hydrogen peroxide (42–45). We detected NOX4 in nuclear, mitochondrial and ER fractions, but only the protein in the ER fraction was depleted upon siRNA transfection (Supplementary Figure 6B). This may reflect that NOX4 protein is more stable in other locations than the ER. To determine if H_2_O_2_ generated by NOX4 is a significant contributor to ROS and quenched by PRDX4, we depleted PRDX4 and NOX4 alone and together and in two cell lines sensitive to PRDX4 depletion and two resistant cell lines. Loss of PRDX4 alone increased the total ROS only in sensitive cells, while no change was seen in resistant cells (Figure 5A and Supplementary Figure 6C). This increase in ROS could be prevented by NOX4 depletion, rendering NOX4 a likely source of the excessive ROS. Knockdown of NOX4 could rescue the inhibition of proliferation caused by loss of PRDX4 while having no effect on proliferation when depleted alone (Figure 5B and Supplementary Figure 6D-E). Finally, NOX4 knockdown also prevented phosphorylation of H2AX caused by loss of PRDX4 (Supplementary Figure 6F). These results demonstrate that NOX4 is the major source of ROS that causes DNA damage and cell toxicity when PRDX4 is depleted.

**Figure 5:**
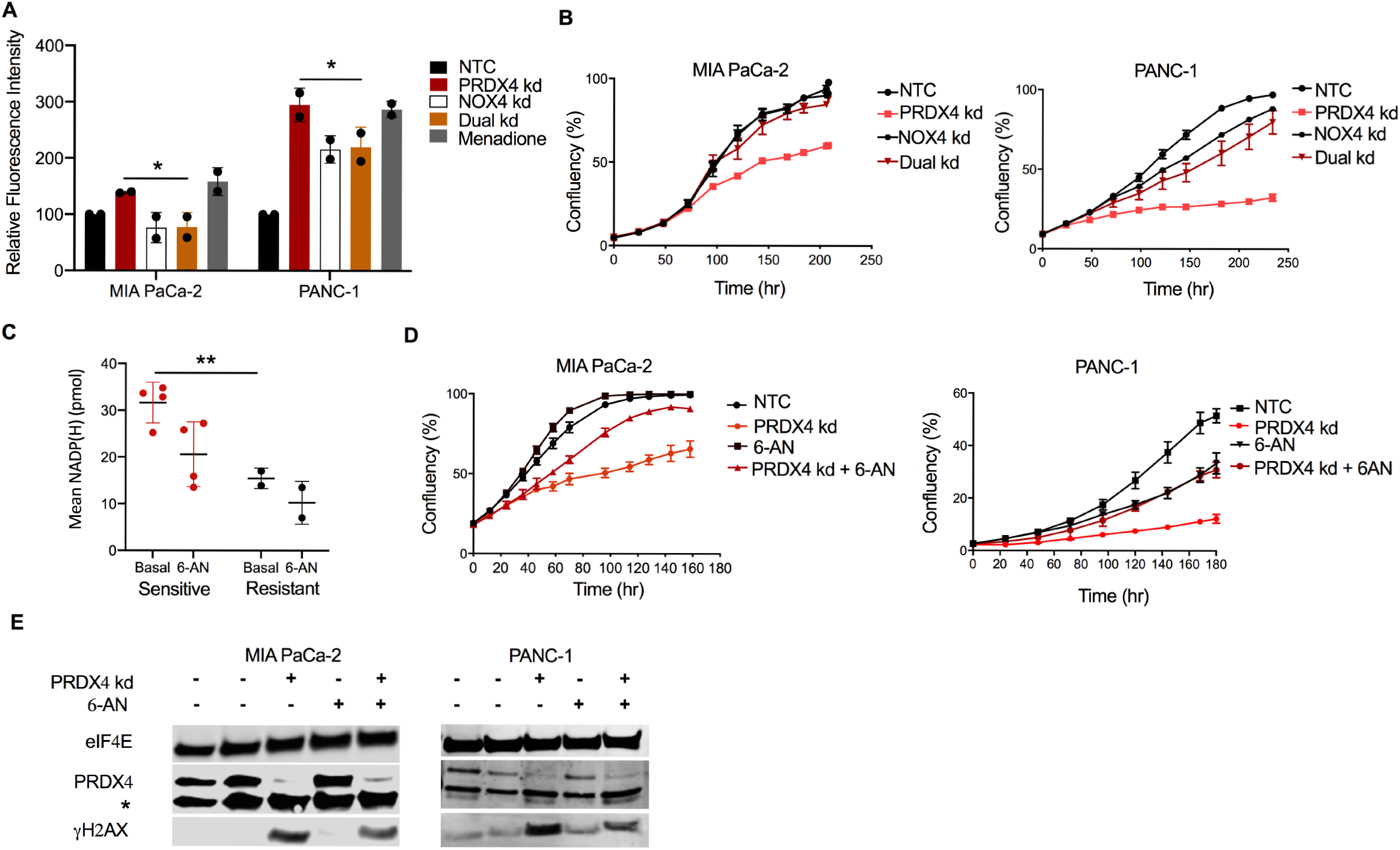
Depletion of NOX4 or NADPH rescues PRDX4 dependency. A) Quantification of relative fluorescence intensity from oxidized CellROX measured by flow cytometry in the indicated cells 72 hours after transfection with siRNA targeting nothing (NTC), PRDX4 (PRDX4 kd), NOX4 (NOX4 kd) or both (Dual kd). Data points represent independent experiments and bars represent the average value ± SD *: p<0.05. B) Cell confluency as a function of time after siRNA transfection as in (A) as measured by automated live cell imaging for the indicated cell lines. Data points represent the average of 3 biological replicates ± SD. C) Quantification of total NADP(H) under basal conditions or after treatment with 5μM 6-aminonicotamide (6-AN) for 24 hours in cells sensitive (MIA PaCa-2, PANC-1, SK-PC-3, BxPC-3) or resistant (Capan-2, GP-3A) to PRDX4 depletion. Each data point represents a cell line. Average value ± SD in each group is indicated. **: p<0.01. D) Cell confluency as a function of time after siRNA transfection as in (A) and/or addition of 5μM 6-AN as measured by automated live cell imaging for the indicated cell lines. Data points represent the average of 3 biological replicates ± SD. E) Western blot showing expression of γH2AX 72 hours after transfection with siRNA as in (A) and/or treatment with 5μM 6-AN from the indicated cell lines. * denotes a cross reacting band.

To assess if basal *NOX4* expression might underlie sensitivity or resistance to PRDX4 depletion, we analyzed RNA sequencing data that accompanied the functional genomic screens. NOX4 mRNA abundance and PRDX4 dependency were uncorrelated (data not shown). Instead, since NOX4 activity relies on the NADPH substrate, we considered whether high NADPH levels might be associated with PRDX4 dependency. Notably, total NADP(H) levels were significantly higher in four cell lines sensitive to PRDX4 depletion compared to two resistant cell lines (p<0.01)(Figure 5C). We manipulated NADPH levels by incubating the cells with 6-aminonicotamide (6-AN), a specific inhibitor of 6-phosphogluconate dehydrogenase (G6PD), which is part of the oxidative arm of the pentose phosphate pathway and a major contributor to cellular NADPH pools. 6-AN reduced NADPH in sensitive cell lines (Figure 5C) and partially rescued the anti-proliferative effects of PRDX4 depletion (Figure 5D). Furthermore, 6-AN also partially reduced H2AX phosphorylation induced by PRDX4 depletion (Figure 5E). Taken together, these data demonstrate that PRDX4 dependency is the result of a metabolic phenotype associated with high NADPH levels in pancreatic cancer cells that drive NOX4-mediated ROS production.

## Discussion

The relationship between oncogenic transformation and redox regulation remains an intense topic of research, but it is increasingly clear that redox vulnerabilities may provide therapeutic opportunities. In pancreatic cancer, oncogenic *KRAS* is known to cause oxidative stress, which stimulates formation of pre-neoplastic lesions and cancer progression (13, 46). At the same time, KRAS has been shown to induce transcription of *NFE2L2* and *GOT1* that contribute to ROS detoxification and tumorigenesis (8, 9). As such, *KRAS* mutant pancreatic cancer cells maintain a delicate balance of ROS to promote proliferation without incurring toxicity. Here we show that dependence on antioxidant proteins of the peroxiredoxin family is a universal feature of pancreatic cancer cell lines (Figure 1A). However, precisely which PRDX family members are essential is highly heterogeneous. Since PRDX family members are expressed in different cellular compartments, this heterogeneity likely reflects the contribution of multiple pathways to the balance between production and detoxification of ROS in specific locations.

Various strategies have been employed in experimental models to exploit the vulnerable redox homeostasis in pancreatic cancer, including disruption of GOT1- and glutamine-dependent NADPH production that provides the reductive power for most cellular anti-oxidant defenses (9, 10, 47). Our results open the possibility of a conceptually orthogonal approach by the exploitation of a NADPH-*driven* redox vulnerability in another cellular compartment. Interestingly, the pathways that support redox homeostasis by high NADPH production in the cytoplasm render the ER fragile due to NOX4-dependent ROS production. The ER is unique in this respect, since other NOX family members primarily produce ROS in the extracellular space. Our results also suggest that flux through the oxidative arm of the pentose phosphate pathway (PPP) contributes significantly to this vulnerability, since the G6PD inhibitor 6-AN could reduce NADPH levels and rescue dependency on PRDX4 (Figure 5C-E). This suggests that the PPP remains active and important in these cells although *KRAS*^G12D^ has been shown to divert significant amounts of glucose intermediates to the non-oxidative arm of the PPP, and to promote alternative NADPH producing pathways (7, 9).

Our data suggest that cell death after PRDX4 knockdown comes as a consequence of H_2_O_2_ generated in the ER diffusing to the nucleus and causing DNA damage (Figure 4). This is consistent with the cellular diffusion distance of H_2_O_2_ (15), and in line with other data demonstrating DNA damage after overexpression of NOX4 (48). Although H_2_O_2_ would be expected to also oxidize proteins, such damage may be more readily reversible, or simply not accumulate to levels that trigger the UPR. It was somewhat surprising that NOX4 knockdown could completely rescue cells from the consequences of PRDX4 depletion, given that ERO1-Lα is considered to be a main generator of hydrogen peroxide in the ER lumen through its activity in disulfide bond formation (49). Furthermore, overexpression or a hyperactive mutant of ERO1-Lα has been shown to affect the oxidation status of PRDX4 (28). Nevertheless, our data are consistent with other published work that also reported an ERO1-Lα-independent source of H_2_O_2_ (50). It is possible that the local organization of oxidases and peroxidases in the ER lumen and membrane contribute to specific interdependencies. Interestingly, human umbilical vein endothelial cells (HUVECs) activate NOX4 upon stimulation with HIV-Tat protein, to produce H_2_O_2_ in the ER lumen and mediate local RAS activation on the ER surface (43). This places NOX4 upstream of wildtype RAS signaling under stress conditions, although other work shows that expression of the NOX4 catalytic subunit *p22*^phox^ is induced by oncogenic KRAS^G12V^ *via* activation of NF-kB (51). In PDAC cells, this stimulated NADH oxidation, glycolysis and proliferation. Clearly, there is more to learn about the interplay between KRAS and NOX4 signaling, and their impact on deregulation of metabolism and linked vulnerabilities in pancreatic cancer.

Regardless, the accompanying DNA damage presents opportunity to combine PRDX4-targeting with other DNA damaging agents such as ionizing radiation, to enhance cell death in a manner specific to the PRDX4-sensitive cancer cells. Recent technological advancements in image-guided radiotherapy has propelled the use of radiotherapy for pancreatic cancer (52). Personalized agents for radiosensitization will be welcomed in this context, and development of PRDX4-targeting agents may be possible. The natural product adenanthin inhibits PRDX1 and 2, but not PRDX4 (53). Adenanthin is thought to interact with Cys173 in PRDX1, which represents the resolving cysteine (C_R)_ important for PRDX reduction and recycling. The structural differences between PRDX family members may provide opportunity for selective activity of inhibitors.

Another important aspect of personalized medicine is patient selection. *PRDX4* is overexpressed in diseases like prostate cancer, lung cancer, colorectal cancer, breast cancer, ovarian cancer and lymph node metastases from oral cavity squamous cell carcinoma (54), and is associated with poor outcome in early stage lung squamous cell carcinoma (55). Here we showed that *PRDX4* expression is associated with aggressive disease, and that higher expression confers poor outcome in resectable pancreatic cancer (Supplementary Figure 2). However, sensitivity to PRDX4 depletion is not a result of “oncogene addiction”, but rather represents a contextual synthetic lethality created by metabolic rewiring. Going forward, NADPH concentration in tumor tissue might serve as a predictive biomarker for sensitivity to PRDX4-targeting. Our data also point to the presence of DNA damage and γH2AX after PRDX4-targeting as a predictor of long-term response.

Our work has focused on pancreatic cancer, and it remains an open question whether targeting PRDX4 may also be relevant for other tumor sites. Others have reported PRDX4 knockdown to hamper proliferation of selected cell lines from lung cancer and GBM (55, 56). It will be important to elucidate the frequency of PRDX4 essentiality in these and other sites, and whether this vulnerability is driven by similar metabolic deregulation.

In conclusion, data presented here highlight the universal reliance of pancreatic cancer cells on ROS detoxification mechanisms, and on PRDX family members in particular, further fueling strategies to target vulnerabilities associated with redox homeostasis. We have uncovered an interesting link between metabolic deregulation and ER oxidative stress mediated by NOX4. We propose that PRDX4 represents a potential molecular target in pancreatic cancer associated with a wide therapeutic window and accompanying predictive biomarkers.

## Materials and methods

### Functional genomics analysis

shRNA Activity Rank Profile (shARP) scores and their standard deviations (SDs) were obtained from a published functional genomics dataset (20). The analysis pipeline was the same as described in Marcotte et al (20) with the following modification. We applied a threshold of approximately 0.4 as defined by the bimodal SD distribution (Supplementary Figure 1A) to define shRNAs with low and high variability. We then used the distributions of shARP scores with low and high variability in order to find cutoffs where the probability of the score belonging to the subpopulation with high SD was higher than the probability of belonging to the subpopulation with low SD (Supplementary Figure 1A). These cutoffs were −0.3 and 0.4, where scores lower than −0.3 reflected a detectable dropout in the screen. As such, essentiality scored on gene level less than −0.3 were used to define essential genes. Analysis was performed in R 3.6.0.

### RNA sequencing of cell lines

Total RNA was extracted and assessed for quality as previously described (57). Sequencing libraries were prepared using either EpiCentre ScriptSeqMR or NuGEN EncoreMR library prep kits following the manufacturer’s instructions. The library size was assessed using a Bioanalyzer DNA 1000 Chip, and the concentration estimated using a Qubit fluorometer and Quant-iT dsDNA BR Assay Kit (Invitrogen). Samples were then loaded on an Illumina HiSeq 2000 sequencer and sequenced using 2×100 paired-end reads. FASTQ files were aligned to genome build hg19 with Gencode v19 gene models using the STAR short-read aligner (v2.3.0). Files containing read counts per Ensembl gene ID, produced by STAR, were merged into a read-count matrix using a bespoke R script, including Ensembl gene annotations. We removed batch effects related to the library preparation kits from the gene expression profiles of 32 sequenced cell lines using ComBat 3.1 (58). To determine the association between essentiality of PRDX1-6 genes and their levels of expression, we obtained R2 and p-values for each gene by fitting linear models. Analysis was performed in R 3.6.0.

### Patient population and RNA sequencing analysis

RNA sequencing data was obtained from patients on the COMPASS trial, as described previously (59). Briefly, biopsies from the primary or metastatic site were collected prior to start of any treatment in a prospective multi-institutional Canadian cohort study. Frozen specimens underwent laser capture microdissection for enrichment of tumor, and Transcript per million (TPM) RNA sequencing data were analyzed as described (59). Modified Moffit subtypes and RNA expression was dichotomized using the maximal chi-squared statistics as described (59).

### Cell culture

PANC-1 (ATCC, CRL-1469), BxPC-3 (ATCC, CRL-1687), SK-PC-3 and GP-9A cells were cultured in RPMI-1640 (Sigma). Capan-2 (ATCC, HTB-80) cells were cultured in McCoy 5A (Sigma). MIA PaCa-2 (ATCC, CRL-11268), GP-2A, GP-3A, GP-5A, GP-9A, GP-10A and GP-16A cells were cultured in DMEM (Sigma). All media was supplemented with 2%-10% fetal bovine serum (Gibco) and cultured under conditions of 5% CO_2_ in air at 37⁰C. For primary pancreatic cancer cells 1X penicillin/streptomycin was added in the growth media. For hypoxia, cells were transferred to a H45 Hypoxystation (Don Whitley Scientific). DNA-PK inhibitor Nu7441 (10 μM, Medchem Express) was added at the time of seeding and replenished one hour before collecting the cells.

### Plasmid construction

Tetracycline/Doxycycline-inducible PRDX4 shRNA oligos (TRCN0000064821 (#4), Broad Institute) were annealed and inserted in the pLKO.1 Tet-On plasmid, a gift from Dmitri Wiederschain (Addgene plasmid # 21915; http://n2t.net/addgene:21915; RRID:Addgene_21915) (60) using the AgeI-HF and EcoRI-HF restriction sites to generate p-lenti-TetOn-SH4. In order to determine the presence of shRNA containing plasmids, sequencing was performed using the primers 5’-GGCAGGGATATTCA and CCATTATCGTTTCAGA-3’.

### Lentiviral production and stable cell line generation

HEK293T cells were transfected with packaging plasmid psPAX2 and envelope plasmid pMD2.G (both gifts from Didier Trono, Addgene plasmid # 12260; http://n2t.net/addgene:12260; RRID:Addgene_12260 and Addgene plasmid # 12259; http://n2t.net/addgene:12259 ; RRID:Addgene_12259) and pLKO.1. Viral supernatant was collected, and titer was calculated using QuickTiter lentivirus Titer Kit (Cell Biolabs, Inc.). Virus was added to MIA PaCa-2 and PANC-1 cells with 10ug/ml polybrene at a MOI of 0.3 and selected with puromycin (3μg/ml) after overnight incubation. Single cells were seeded and propagated to create stable clones. For subsequent experiments, doxycycline (2μg/ml) was added at the time of cell seeding.

### siRNA transfections

PRDX4 siRNA (HSS173720, Life Tech), NOX4 siRNA (HSS121312, Life Tech), ERO1lα siRNA (HSS121196) or non-targeted control (NTC) siRNA (Thermofisher Scientific) were transfected using the lipofectamine RNAiMAX reverse transfection method (Thermofisher Sientific) at final concentration of 2.5nM.

### Cell proliferation, viability and clonogenicity

siRNA or compounds were added at time of seeding and multi well plates placed in the IncuCyte ZOOM Kinetic Imaging System for monitoring cell confluency. For all cell lines, it was verified that they grew in a monolayer and that confluence was proportional to cell number. Alamar blue was added to the media for viability measurements and fluorescence was read at 590nm using the OMEGA plate reader (BMG Labtech) after 3 hours. For clonogenic survival, single cells were plated in 6-cm dishes and incubated for 14 days. Bromophenol blue was used to stain colonies. Plating efficiency (PE) was calculated as # colonies (>50 cells) divided by # cells seeded. Surviving fraction was calculated as PE(treatment)/PE(control).

### Xenograft establishment and treatments

All animal experiments were performed according to operating procedures approved by the Princess Margaret Cancer Centre Animal Care Committee, aligned with guidelines from the Canadian Council on Animal Care. For subcutaneous tumors, female NOD-Rag1^null^/IL2rg^null^ (NRG) mice (Jax/7799) bred in house were implanted with 4×10^6^ MIA PaCa-2 or PANC-1 cells in the flank or shoulder. Once tumors reached 200mm^3^ doxycyclin (625 mg/kg) was administered in chow. Tumors were measured twice a week by caliper until they exceeded 800mm^3^, upon which the mice were sacrificed, and tumors excised and frozen. Radiation therapy started one week after Dox treatment with 5 doses of 7 Gy given over 10 days. Anesthetized mice were immobilized for cone-beam CT imaging for planning and subsequent irradiation using an X-Rad 225Cx (Precision X-ray) at a dose rate of 3 Gy/minute. For orthotopic tumors, female NRG mice were first implanted with 4×10^6^ MIA PaCa-2 or PANC-1 cells in the flank to establish donor tumors. Once tumors reached 800mm^3^ they were excised and 1mm by 1mm pieces were sutured on the pancreas tail of acceptor mice. 1 week after tumor implantation, doxycycline (625mg/kg) was administered in the chow. 3 days thereafter, weekly magnetic resonance imaging (MRI) using 7T MRI system (Bruker) was used to measure tumor growth. Overall survival and disease free survival analyses were performed by fitting a Cox proportional hazards regression model using "survival" package in R (61, 62) in R 3.6.0. P-values were obtained using the likelihood ratio tests.

### RT-qPCR

mRNA was extracted according to the PureLink RNA Mini Kit (Life Tech) protocol and quantified with the NanoDrop lite spectrophotometer (Thermofisher Scientific). Reverse transcription was performed according to the qScript cDNA SuperMix (Quanta Biosciences) protocol. Quantitative PCR analysis was performed using SYBR green (perfecta SYBR Green Supermix, Quanta Biosciences) and primers listed in Supplementary Table 1.

### Immunoblotting

Subcellular fractionation was carried out as descried earlier (63). Otherwise, protein was extracted in RIPA lysis buffer (Norgen Biotek Corp.) supplemented with protease and phosphatase inhibitor cocktail (Thermofisher Scientific) and EDTA. Protein quantification was performed according to the bicinchoninic acid method (Pierce BCA Protein Assay Kit, Life Tech) using the OMEGA plate reader (BMG Labtech) at 562nm. Proteins were resolved by SDS-PAGE and proteins were transferred onto a PVDF membrane. The membranes were incubated with primary antibodies listed in Supplementary Table 2 overnight at 4^°^C. IR-conjugated secondary antibodies (all Mandel) were applied for 1 h at room temperature and included: IRDye^®^ 800CW Donkey anti-Rabbit, IRDye^®^ 800CW Donkey anti-Mouse, IRDye^®^ 680RD Donkey anti-Rabbit, and IRDye^®^ 680CW Goat anti-Mouse. Proteins were visualized on the near infrared Odyssey Li-Cor (LI-COR Biosciences), followed with data quantification using Image Studio 5.02 Lite.

### Immunofluorescence

Cells were seeded in 8 well chamber slides, fixed in paraformaldehyde for 20 minutes at room temperature and permeabilized using 0.5% triton-X 100 in PBS for 10 minutes. Blocking was done using 5% bovine serum albumin (BSA) in PBS for 1 hour. Mouse anti-γH2AX (1:100 in 2% BSA in PBS) primary antibody was applied for 1 hour and Anti-Rabbit IgG Alexa Fluor 488 (1:200 in 2% BSA in PBS, 4408S, New England biolabs) was applied for 1 hour. Nuclei were stained with 4-6-diamidino-2-phenylindole (DAPI) for 5 minutes. All washing was performed with 0.025% tween20 in PBS. Slides were mounted with Vectashield antifade mounting media (Vector laboratories) and a coverslip before imaging. Imaging for fluorescence was done using ZEISS LSM 710 Confocal microscope (C-Apo63x/1.4 NA objective, oil), followed by image analysis using ZEN image analysis v2.1 and Fiji.

### Comet Assay

Cells were collected, counted, mixed in low melting agarose (1%) at a concentration of 40,000/ml and applied to slides. Electrophoresis was performed under both alkaline and neutral buffer conditions as described in the CometAssay Kit (Trevigen). The slides were dried at 37^°^C and stained with SYBR gold nucleic acid gel stain (Thermofisher Scientific). Slides were mounted with Vectashield antifade mounting media (Vector laboratories) and a coverslip before imaging with a Zeiss AxioObserver microscope (20X objective, Olympus). Analysis was done using ImageJ software.

### Flow Cytometry

CellROX deep-red (Thermofisher Scientific) or propidium iodide (Abcam) was applied to cells according to manufacturer’s protocol. Flow cytometry was performed in a LSM-OICR (Becton Dickinson) and analysed using FlowJo software version 10 (FlowJo LLC.).

### Statistical analysis

Unless otherwise specified, data were analyzed with the use of Graph Pad Prism 5 (Graph Pad Software Inc., San Diego, CA, USA) with either unpaired students t-test or one-way ANOVA with Bonferroni correction.

## Author contributions

P.J and M.K. designed the study. P.J., E.M., M.X., F.J., P.D. and S.C. conducted and analyzed laboratory experiments, A.D-G., K.R.B. and G.H.J. analyzed functional genomics, RNAseq and clinical data, F.N., J.M., D.H., P.C.B., B.G.W and M.K. provided resources and supervised analysis, P.J. and M.K. wrote the manuscript, all authors edited the manuscript.

## Acknowledgements

We appreciate the advice of Azin Sayad on analysis of the functional genomics dataset. Funding for this work was provided by the Cancer Research Society / Robert Lutterman Memorial Fund (Operating Grant to M.K., D.H., P.C.B.), The Terry Fox Foundation/Research Institute (New Frontiers Research Program PPG09-020005 to M.K., B.G.W., D.H.), The Princess Margaret Cancer Foundation (Invest in Research award to M.K. and Postdoctoral Fellowship to P.J.), the Strategic Training in Translational Radiation Medicine for the 21^st^ Century (STARS21) Program (Fellowship to P.J.), the NIH/NCI (P30CA016042 to P.C.B.), the Ontario Ministry of Health and Princess Margaret Cancer Centre.

---

**Figure S1:**
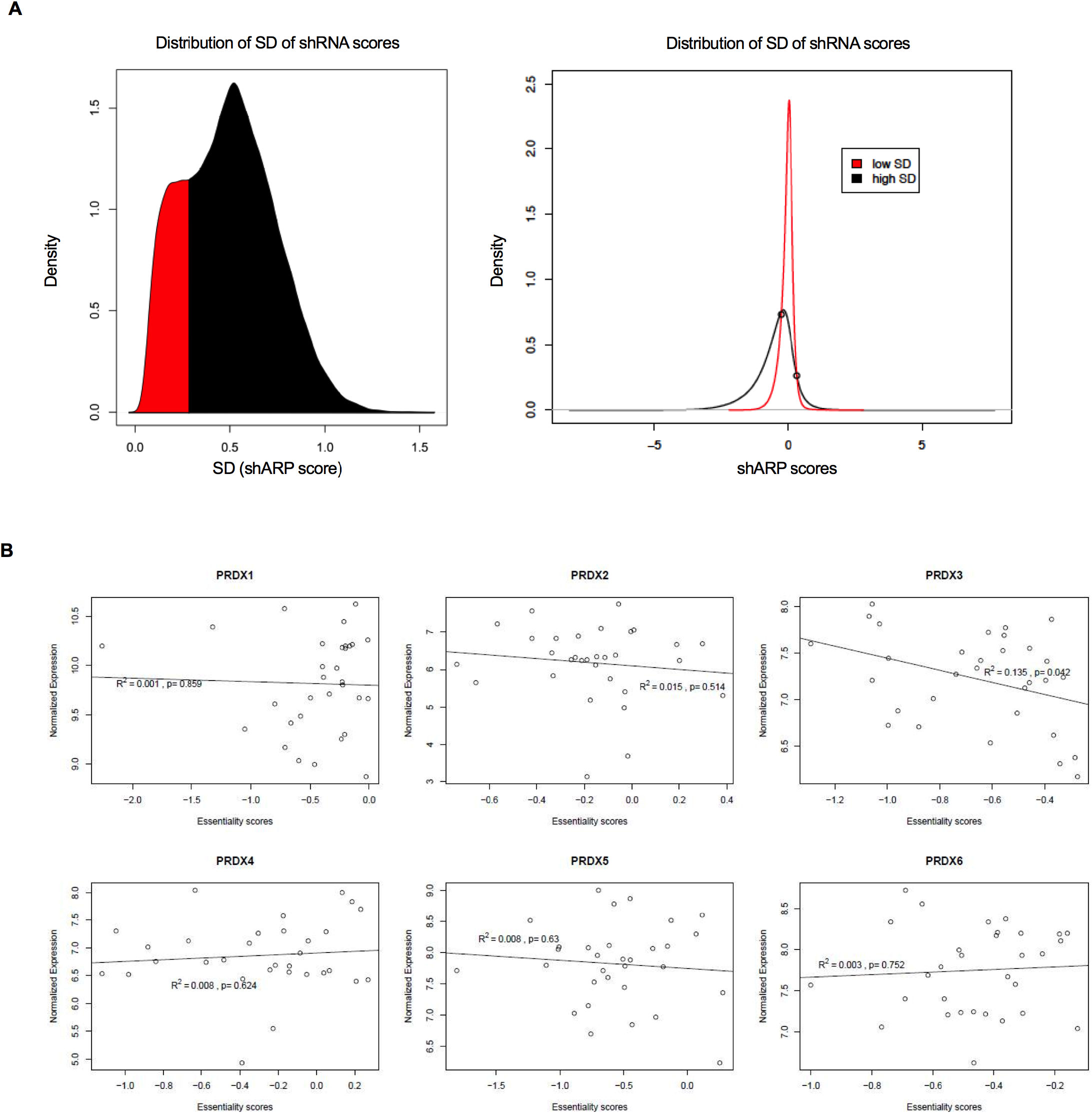
A) *Leftpanel:* Density plot showing distribution of Standard Deviations (SD) of shRNA Activity Rank Profile scores (shARP; (20)) for all genes pooled from all samples. The distribution exhibits 2 subpopulations: low SD and high SD, shown in red and in black, respectively. *Right panel:* Density plots showing distributions of shARP scores the low SD and the high SD subpopulations (shown in red and black subpopulations, respectively). Thresholds marked by black circles were set at score values at which the probability of a score belonging to a specific distribution (high or low SD) switches. shARP scores in below the lower threshold (black circle on the left) were considered to represent essentiality. B) Association between essentiality scores on gene level and mRNA expression levels of *PRDXJ-6* genes. R2 and p-values for each gene were obtained by fitting linear models.

**Figure S2:**
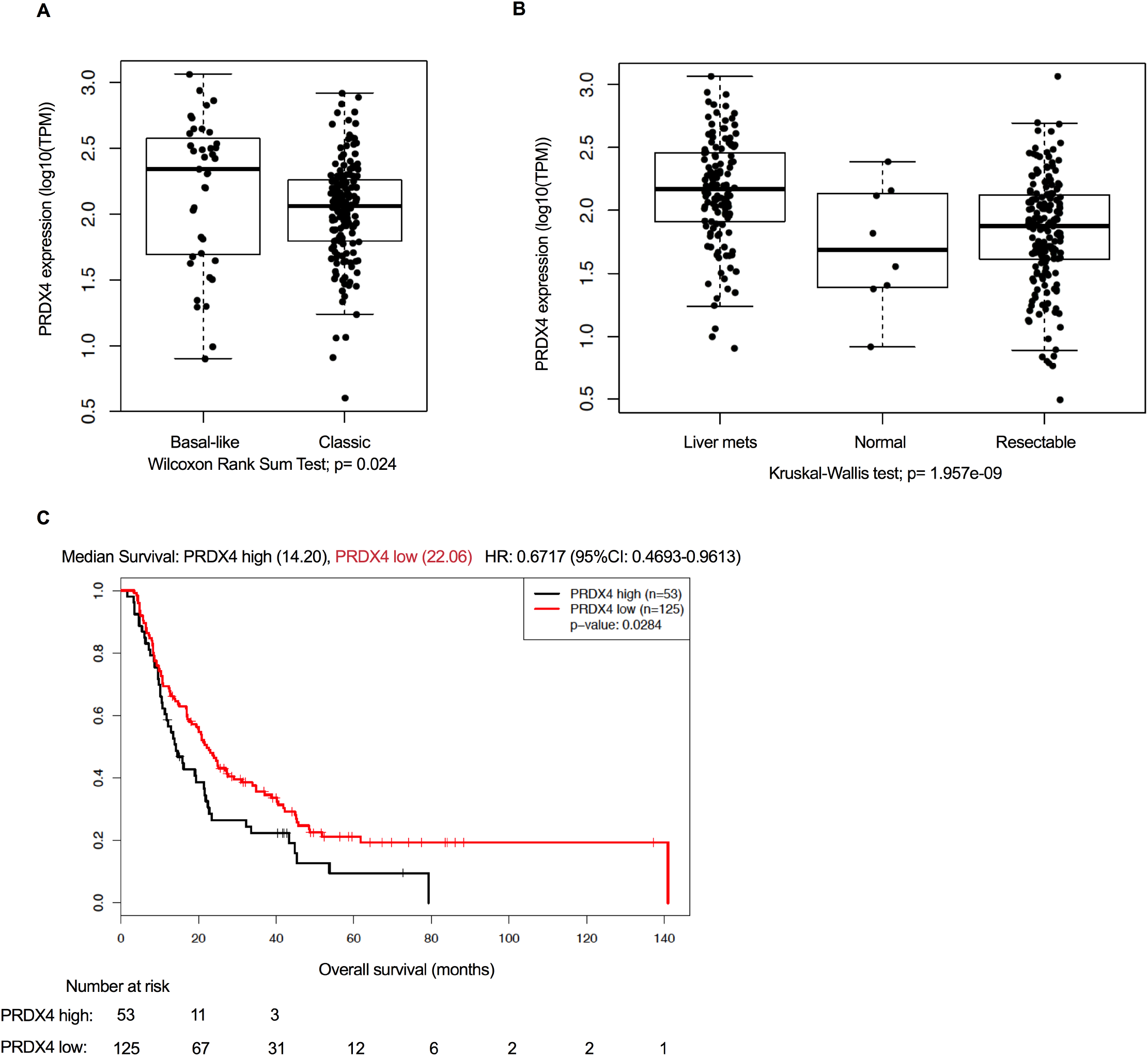
A) PRDX4 expression by RNA sequencing from 195 advanced pancreatic cancer biopsies (29,59) versus modified Moffit subtypes (30). B) PRDX4 expression by RNA sequencing in biopsies from 182 resectable localized pancreatic cancers, 8 adjacent normal tissues and 128 pancreatic cancer liver metastases. C) Kaplan-Meier curve of Overall Survival in 178 patients with resectable pancreatic cancer stratified by median PRDX4 expression.

**Figure S3:**
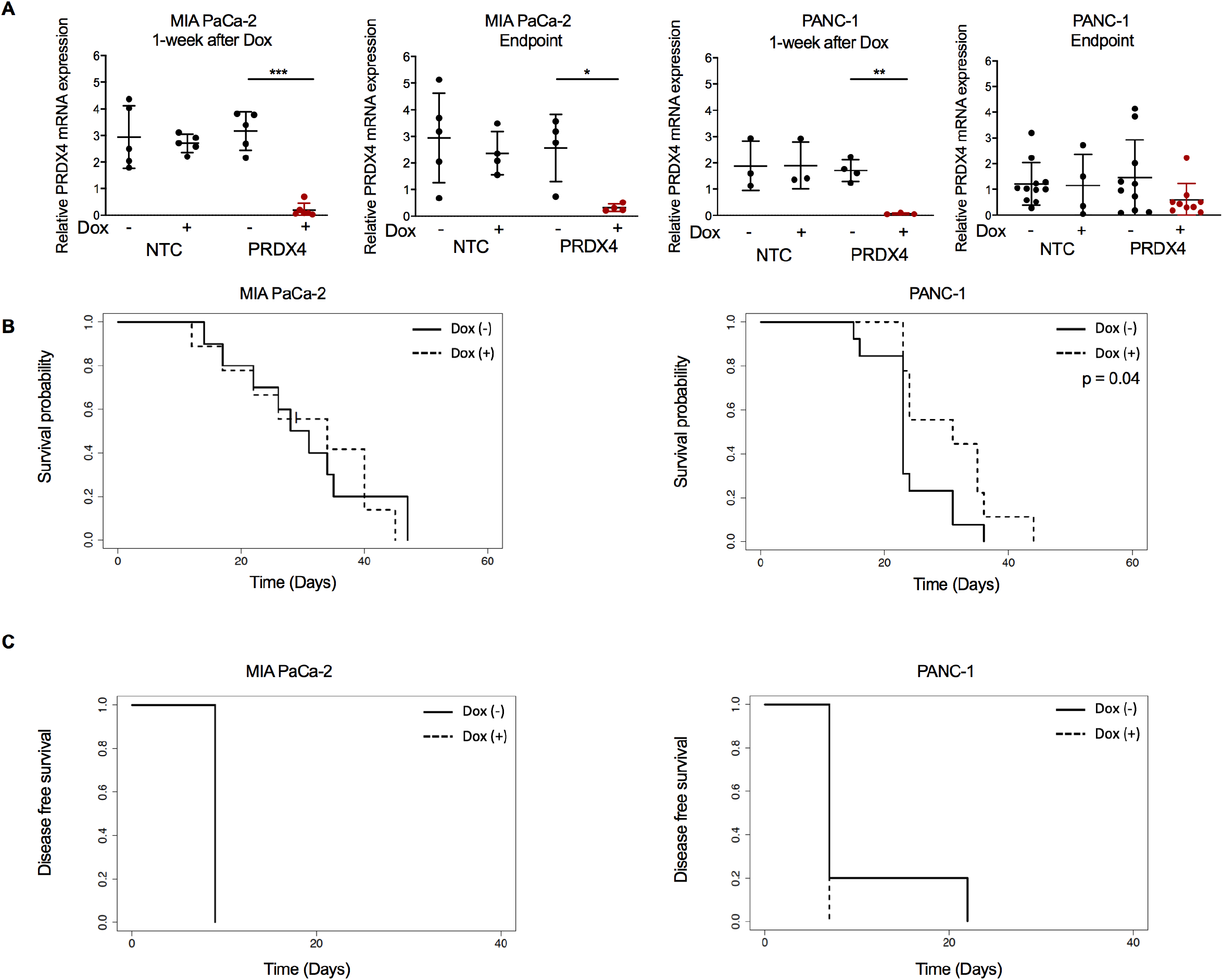
A) Quantification ofPRDX4 mRNA expression in subcutaneous tumors established from MIA PaCa-2 or PANC-1 cells expressing Dox-inducible shRNA targeting nothing (NTC) or PRDX4. Dox was administered in the chow when tumors reached 200mm^3^, and RNA was extracted after one week or at endpoint (>800mm^3^). Data points represent individual tumors; the average value is indicated± SD. Relative expression is normalized to expression of HPRTl in each sample.***: p<0.001;**: p<0.01;*: p<0.05. B) Survival probability ofmice carrying subcutaneous MIA PaCa-2 or PANC-1 tumors expressing Dox-inducible shRNA targeting nothing (NTC) as a function of time after Dox administration in the chow. C) Disease free survival of mice carrying orthotopic MIA PaCa-2 or PANC-1 tumors expressing Dox-inducible shRNA targeting nothing (NTC) as a function of time after Dox administration in the chow.

**Figure S4:**
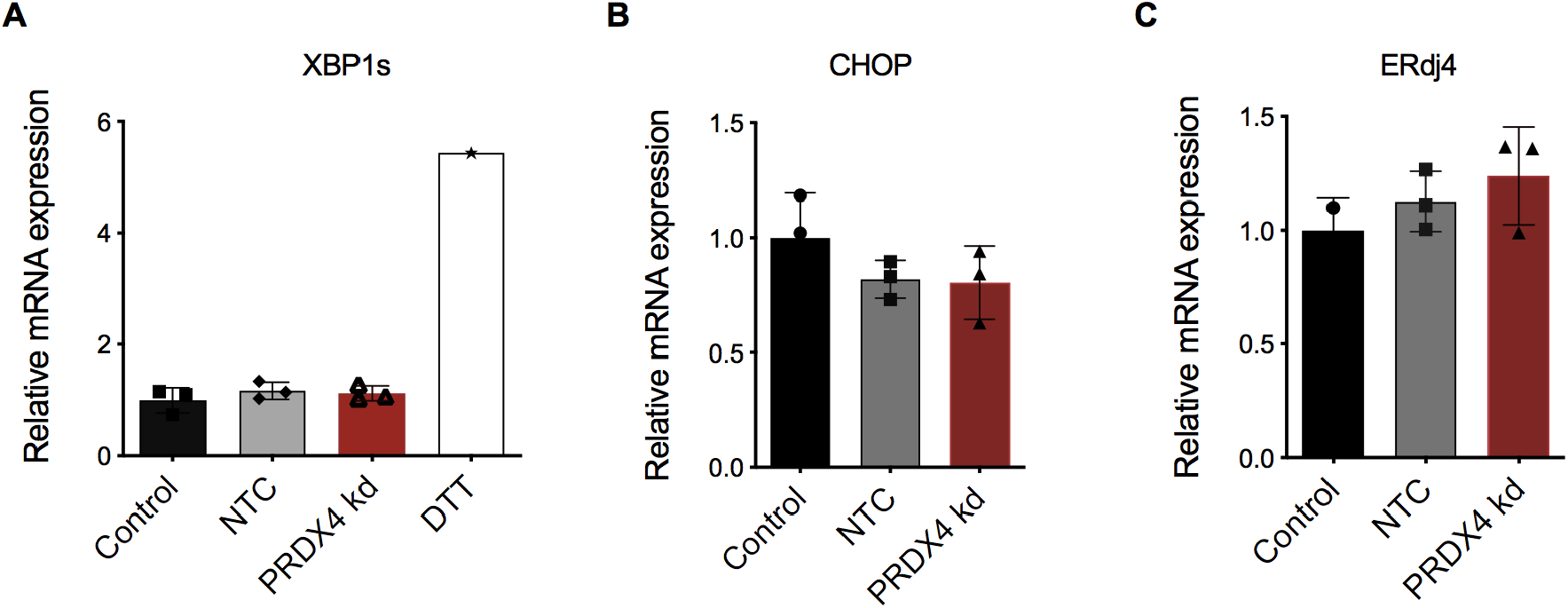
Quantification of relative mRNA expression of A) spliced XBPl (XBPls), B) CHOP or C) ERdj4 from PANC-1 cells 72 hours after transfection with siRNA targeting nothing (NTC) or PRDX4 (PRDX4 kd) or 1.5 hours after exposure to 5mM DTT. Data points represent technical replicates, the bar represents the average value ± SD. Relative expression is normalized to expression ofHPRTl and to the untreated sample.

**Figure S5:**
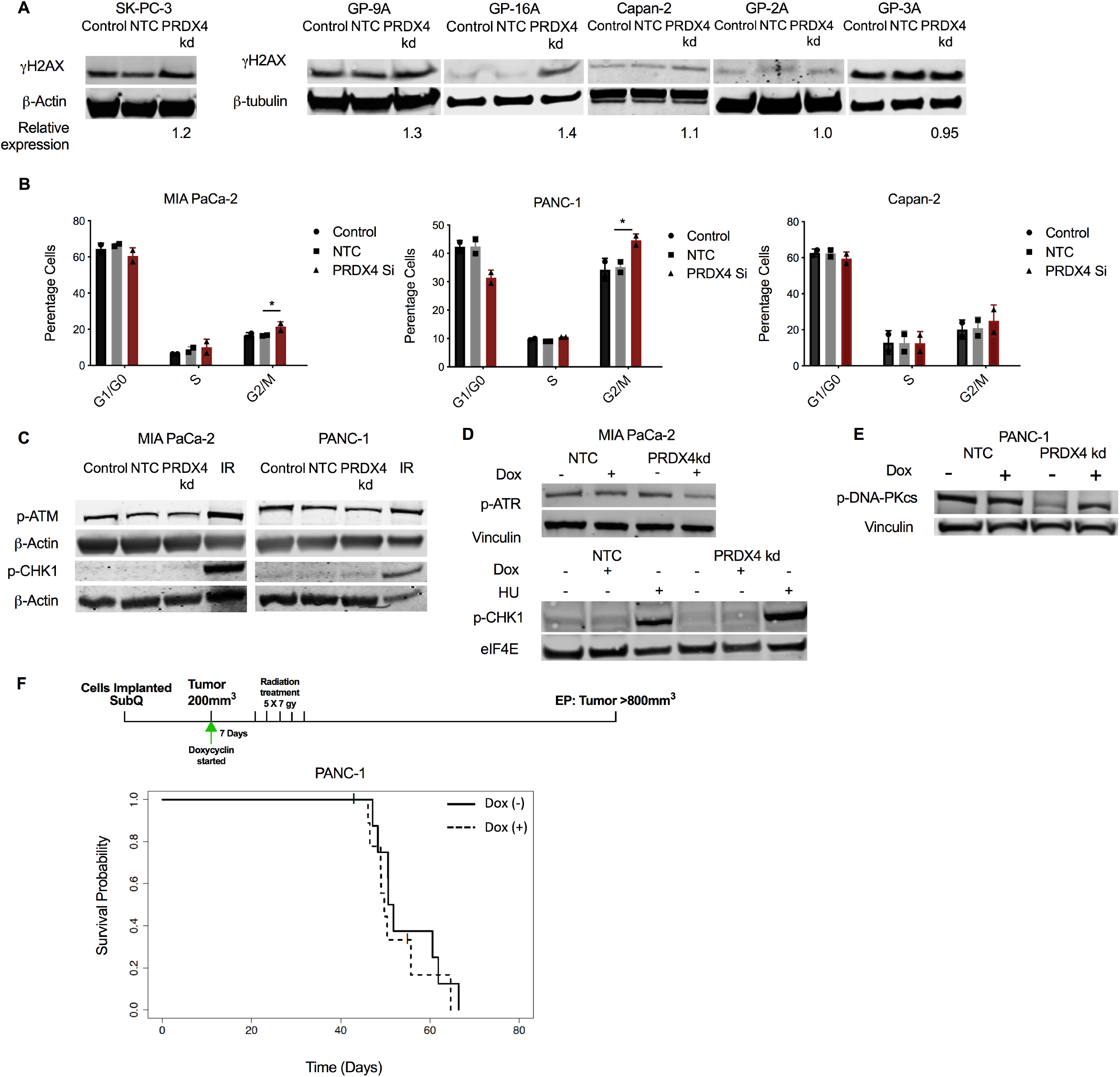
A) Western blot showing expression ofyH2AX in 3 cell lines sensitive (SK-PC-3, GP-9A, GP-16A) and 3 cell lines resistant (Capan-2, GP-3A, GP-2A) to PRDX4 depletion, 72 hours after transfection with siRNA targeting nothing (NTC) or PRDX4 (PRDX4 kd). (The P-Tubulin loading controls are the same as Figure lBii.). Relative expression ofyH2AX after PRDX4 kd was normalised to NTC in each cell line. B) Quantification ofpercentage ofcells from indicated cell lines in indicated phases of the cell cycle as measured by flow cytometry 72 hours after transfection as in (A). Each data point represents a biological replicate, bars represent the average value± SD. *: p<0.05. C) Western blot showing expression ofp-ATM and p-CHKl in MIA PaCa-2 and PANC-1 cells after transfection as in (A) or 30 minutes after 15 Gy ofionizing radiation (IR). D) Western blot showing expression ofp-ATM and p-CHKl in MIA PaCa-2 cells expressing Dox-inducible shRNA targeting nothing (NTC) or PRDX4 (PRDX4 kd) 72 hours after administration ofDox or 5μM hydroxyurea (HU) (24 hours). (The e1F4E loading control is the same as in Figure 4G.) E) Western blot showing expression ofp-DNA-PKcs and yH2AX in PANC-1 cells expressing Dox-inducible shRNA targeting nothing (NTC) or PRDX4 (PRDX4 kd) 72 hours after administration of Dox. F) Survival probability of mice carrying subcutaneous PANC-1 tumors expressing Dox-inducible shRNA targeting nothing (NTC) as a function of time after Dox administration in the chow. Tumors were irradiated with 5 times 7Gy over 10 days starting on Day 7.

**Figure S6:**
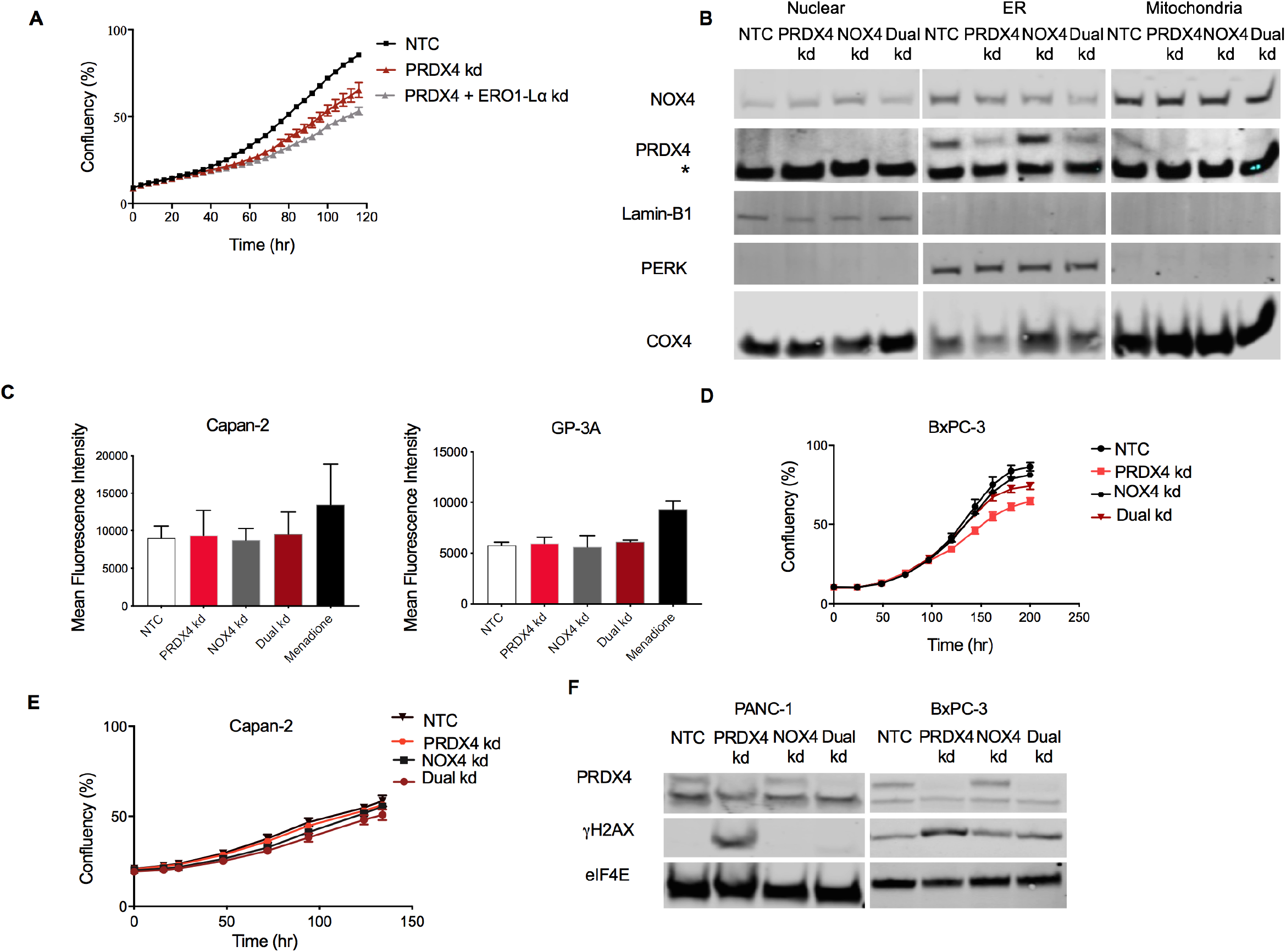
PANC-1 cell confluency as a function of time after transfection with siRNA targeting nothing (NTC), PRDX4 (PRDX4 kd) or PRDX4 together with EROl-La (PRDX4 + EROl-La kd) as measured by automated live cell imaging. Data points represent the average of 3 biological replicates± SD. B) Western blot showing expression ofNOX4, PRDX4, Lamin-Bl (Nuclear marker), PERK (ER marker) and COX4 (Mitochondria marker) after subcellular fractionation of BxPC-3 cells 72 hours after transfection with siRNA targeting nothing (NTC), PRDX4 (PRDX kd), NOX4 (NOX4 kd) or both (Dual kd). C) Quantification of mean fluorescence intensity of oxidized CellROX measured by flow cytometry from two cell lines resistant to PRDX4 depletion 72 hours following transfection with siRNA as indicated, or exposure to 100 μM menadione for 1 hour. BxPC-3 (D) and Capan-2 (E) cell confluency as a function of time after siRNA transfection with siRNA targeting nothing (NTC), PRDX4 (PRDX4 kd), NOX4 (NOX4 kd) or both (Dual kd) as measured by automated live cell imaging. Data points represent the average of 3 biological replicates± SD. F) Western blot showing expression of yH2AX in PANC-1 and BxPC-3 cells 72 hours after siRNA transfection as before.

**Supplementary Table 1:**
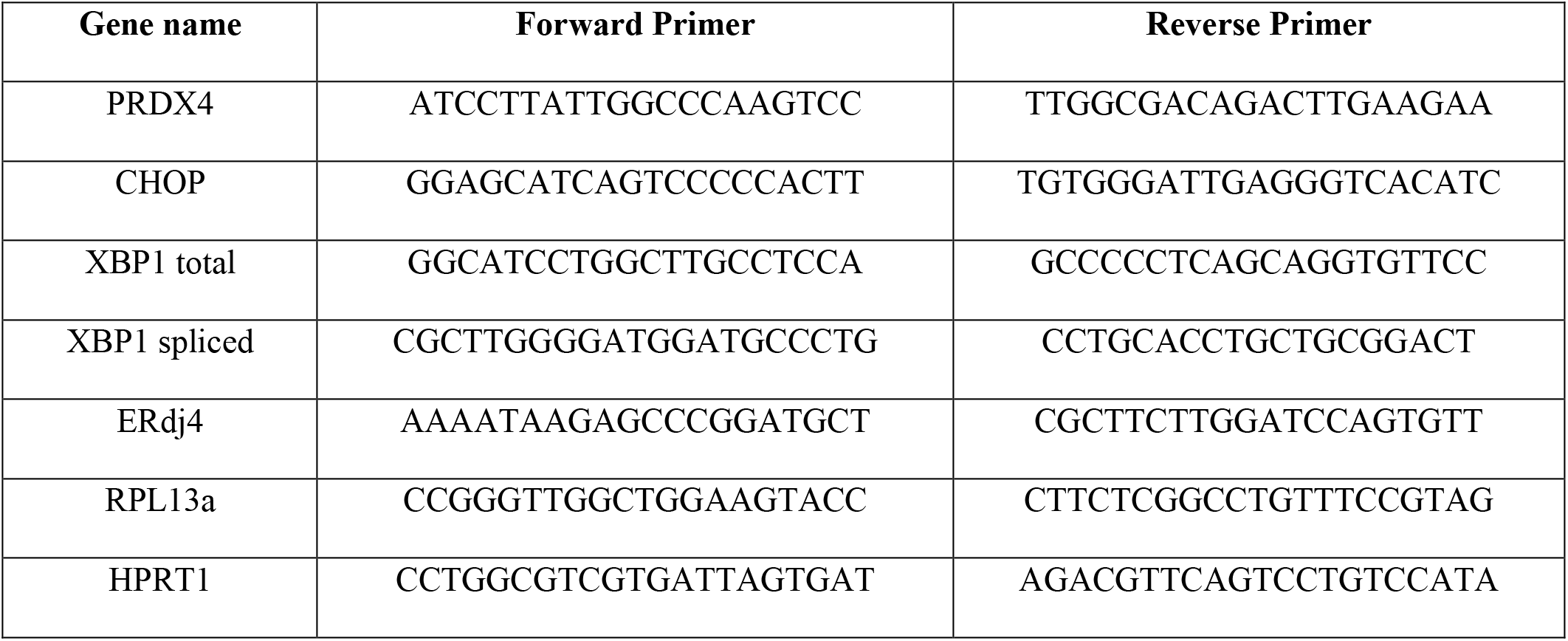
qPCR primers

**Supplementary table 2:**
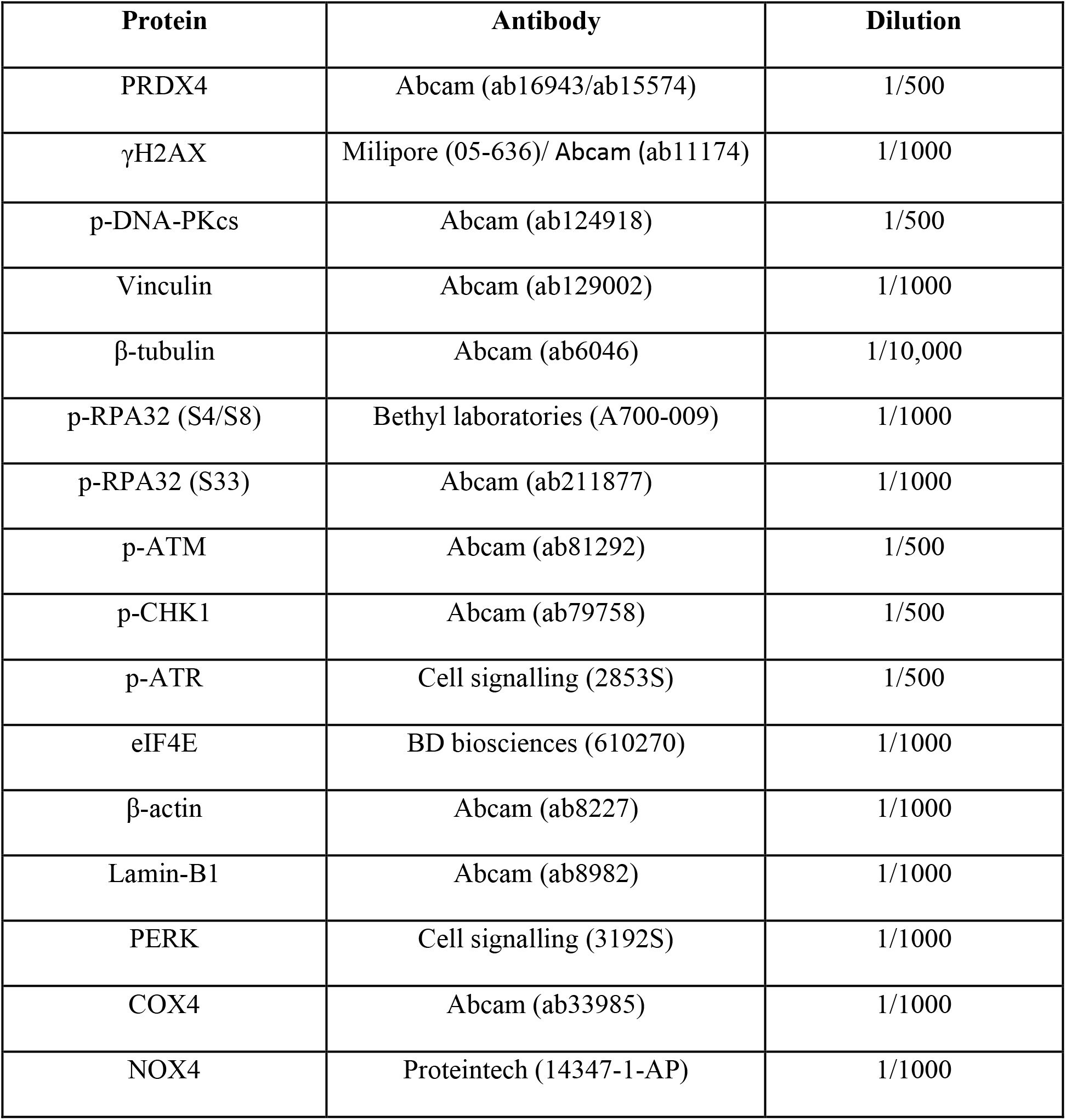
Antibodies

## REFERENCES

1. Collisson EA, Bailey P, Chang DK, Biankin, AV. Molecular subtypes of pancreatic cancer. Nature Reviews Gastroenterology & Hepatology. 2019;16(4):207–220

2. Bailey P, et al. Genomic analyses identify molecular subtypes of pancreatic cancer. Nature. 2016;531(7592):47–52

3. Cortés-Ciriano I, et al. Comprehensive analysis of chromothripsis in 2,658 human cancers using whole-genome sequencing. Nature genetics. 2020/03// 2020;52(3):331–341 doi:10.1038/s41588-019-0576-7

4. Notta F, et al. A renewed model of pancreatic cancer evolution based on genomic rearrangement patterns. Nature. 2016;538(7625):378–382 doi:10.1038/nature19823

5. Canon J, et al. The clinical KRAS(G12C) inhibitor AMG 510 drives anti-tumour immunity. Nature. Nov 2019;575(7781):217–223 doi:10.1038/s41586-019-1694-1

6. Hayes JD, Dinkova-Kostova AT, Tew, KD. Oxidative Stress in Cancer. Cancer Cell. 2020;

7. Ying H, et al. Oncogenic Kras maintains pancreatic tumors through regulation of anabolic glucose metabolism. Cell. 2012;149(3):656–670

8. DeNicola GM, et al. Oncogene-induced Nrf2 transcription promotes ROS detoxification and tumorigenesis. Nature. Jul 6 2011;475(7354):106–9 doi:10.1038/nature10189

9. Son J, et al. Glutamine supports pancreatic cancer growth through a KRAS-regulated metabolic pathway. Nature. 2013;496(7443):101–105

10. Chakrabarti G, et al. Targeting glutamine metabolism sensitizes pancreatic cancer to PARP-driven metabolic catastrophe induced by ß-lapachone. Cancer Metab. 2015;3:12. doi:10.1186/s40170-015-0137-1

11. Storz P. KRas, ROS and the initiation of pancreatic cancer. Small GTPases. 2017;8(1):38–42

12. Weinberg F, et al. Mitochondrial metabolism and ROS generation are essential for Kras-mediated tumorigenicity. Proceedings of the National Academy of Sciences. 2010;107(19):8788–8793

13. Liou GY, et al. Mutant KRas-Induced Mitochondrial Oxidative Stress in Acinar Cells Upregulates EGFR Signaling to Drive Formation of Pancreatic Precancerous Lesions. Cell reports. Mar 15 2016;14(10):2325–36 doi:10.1016/j.celrep.2016.02.029

14. Schieber M, Chandel, NS. ROS function in redox signaling and oxidative stress. Current biology. 2014;24(10):R453–R462.

15. Gorrini C, Harris IS, Mak, TW. Modulation of oxidative stress as an anticancer strategy. Nat Rev Drug Discov. Dec 2013;12(12):931–47 doi:10.1038/nrd4002

16. Martin RE, Cao Z, Bulleid, NJ. Regulating the level of intracellular hydrogen peroxide: the role of peroxiredoxin IV. Portland Press Ltd.; 2014.

17. Perkins A, Nelson KJ, Parsonage D, Poole LB, Karplus, PA. Peroxiredoxins: guardians against oxidative stress and modulators of peroxide signaling. Trends in biochemical sciences. 2015;40(8):435–445

18. Rhee SG, et al. Intracellular messenger function of hydrogen peroxide and its regulation by peroxiredoxins. Current opinion in cell biology. 2005;17(2):183–189

19. Wong C-M, et al. Characterization of human and mouse peroxiredoxin IV: evidence for inhibition by Prx-IV of epidermal growth factor-and p53-induced reactive oxygen species. Antioxidants and Redox Signaling. 2000;2(3):507–518

20. Marcotte R, et al. Essential gene profiles in breast, pancreatic, and ovarian cancer cells. Cancer discovery. 2012;2(2):172–189

21. Zhang Y-G, et al. Featured article: accelerated decline of physical strength in peroxiredoxin-3 knockout mice. Experimental Biology and Medicine. 2016;241(13):1395–1400

22. Neumann CA, et al. Essential role for the peroxiredoxin Prdx1 in erythrocyte antioxidant defence and tumour suppression. Nature. Jul 31 2003;424(6948):561–5 doi:10.1038/nature01819

23. Lee TH, et al. Peroxiredoxin II is essential for sustaining life span of erythrocytes in mice. Blood. Jun 15 2003;101(12):5033–8 doi:10.1182/blood-2002-08-2548

24. Argyropoulou V, et al. Peroxiredoxin-5 as a Novel Actor in Inflammation and Tumor Suppression. Free Radical Biology and Medicine. 2016;100:S92.

25. Wang X, et al. Mice with targeted mutation of peroxiredoxin 6 develop normally but are susceptible to oxidative stress. J Biol Chem. Jul 4 2003;278(27):25179–90 doi:10.1074/jbc.M302706200

26. Iuchi Y, et al. Peroxiredoxin 4 knockout results in elevated spermatogenic cell death via oxidative stress. Biochem J. Apr 1 2009;419(1):149–58 doi:10.1042/bj20081526

27. Zito E. PRDX4, an endoplasmic reticulum-localized peroxiredoxin at the crossroads between enzymatic oxidative protein folding and nonenzymatic protein oxidation. Antioxidants & redox signaling. 2013;18(13):1666–1674

28. Tavender TJ, Bulleid, NJ. Peroxiredoxin IV protects cells from oxidative stress by removing H_2_O_2_ produced during disulphide formation. Journal of cell science. 2010;123(15):2672–2679

29. Aung KL, et al. Genomics-Driven Precision Medicine for Advanced Pancreatic Cancer: Early Results from the COMPASS Trial. Clin Cancer Res. Mar 15 2018;24(6):1344–1354 doi:10.1158/1078-0432.Ccr-17-2994

30. Moffitt RA, et al. Virtual microdissection identifies distinct tumor- and stroma-specific subtypes of pancreatic ductal adenocarcinoma. Nat Genet. Oct 2015;47(10):1168–78 doi:10.1038/ng.3398

31. Mathieson T, et al. Systematic analysis of protein turnover in primary cells. Nature Communications. 2018/02/15 2018;9(1):689. doi:10.1038/s41467-018-03106-1

32. Wilson WR, Hay, MP. Targeting hypoxia in cancer therapy. Nature Reviews Cancer. 2011;11(6):393–410

33. Edmondson R, Broglie JJ, Adcock AF, Yang, L. Three-dimensional cell culture systems and their applications in drug discovery and cell-based biosensors. Assay and drug development technologies. 2014;12(4):207–218

34. Huynh AS, et al. Development of an orthotopic human pancreatic cancer xenograft model using ultrasound guided injection of cells. PLoS One. 2011;6(5):e20330. doi:10.1371/journal.pone.0020330

35. Zito E. ERO1: A protein disulfide oxidase and H_2_O_2_ producer. Free Radic Biol Med. Jun 2015;83:299–304. doi:10.1016/j.freeradbiomed.2015.01.011

36. Wang M, Kaufman, RJ. The impact of the endoplasmic reticulum protein-folding environment on cancer development. Nature Reviews Cancer. 2014;14(9):581–597

37. Fernandez-Capetillo O, et al. DNA damage-induced G 2–M checkpoint activation by histone H2AX and 53BP1. Nature cell biology. 2002;4(12):993–997

38. Olive PL, Johnston PJ, Banath JP, Durand, RE. The comet assay: A new method to examine heterogeneity associated with solid tumors. Nature Medicine. 1998/01/01 1998;4(1):103–105 doi:10.1038/nm0198-103

39. Liu S, et al. Distinct roles for DNA-PK, ATM and ATR in RPA phosphorylation and checkpoint activation in response to replication stress. Nucleic Acids Res. Nov 2012;40(21):10780–94 doi:10.1093/nar/gks849

40. Ying S, et al. DNA-PKcs and PARP1 Bind to Unresected Stalled DNA Replication Forks Where They Recruit XRCC1 to Mediate Repair. Cancer Res. Mar 1 2016;76(5):1078–88 doi:10.1158/0008-5472.Can-15-0608

41. Gross E, et al. Generating disulfides enzymatically: reaction products and electron acceptors of the endoplasmic reticulum thiol oxidase Ero1p. Proceedings of the National Academy of Sciences. 2006;103(2):299–304

42. Bedard K, Krause, KH. The NOX family of ROS-generating NADPH oxidases: physiology and pathophysiology. Physiological reviews. Jan 2007;87(1):245–313 doi:10.1152/physrev.00044.2005

43. Wu RF, Ma Z, Liu Z, Terada, LS. Nox4-derived H_2_O_2_ mediates endoplasmic reticulum signaling through local Ras activation. Mol Cell Biol. Jul 2010;30(14):3553–68 doi:10.1128/MCB.01445-09

44. Pendyala S, Natarajan, V. Redox regulation of Nox proteins. Respir Physiol Neurobiol. Dec 31 2010;174(3):265–71 doi:10.1016/j.resp.2010.09.016

45. Shanmugasundaram K, et al. NOX4 functions as a mitochondrial energetic sensor coupling cancer metabolic reprogramming to drug resistance. Nature communications. 2017;8(1):1–16

46. Cheung EC, et al. Dynamic ROS Control by TIGAR Regulates the Initiation and Progression of Pancreatic Cancer. Cancer Cell. Feb 10 2020;37(2):168–182e4. doi:10.1016/j.ccell.2019.12.012

47. Yoshida T, et al. A covalent small molecule inhibitor of glutamate-oxaloacetate transaminase 1 impairs pancreatic cancer growth. Biochem Biophys Res Commun. Feb 12 2020;522(3):633–638 doi:10.1016/j.bbrc.2019.11.130

48. Weyemi U, et al. ROS-generating NADPH oxidase NOX4 is a critical mediator in oncogenic H-Ras-induced DNA damage and subsequent senescence. Oncogene. 2012;31(9):1117–1129

49. Enyedi B, Varnai P, Geiszt, M. Redox state of the endoplasmic reticulum is controlled by Ero1L-alpha and intraluminal calcium. Antioxid Redox Signal. Sep 15 2010;13(6):721–9 doi:10.1089/ars.2009.2880

50. Konno T, et al. ERO1-independent production of H_2_O_2_ within the endoplasmic reticulum fuels Prdx4-mediated oxidative protein folding. J Cell Biol. Oct 26 2015;211(2):253–9 doi:10.1083/jcb.201506123

51. Ju HQ, et al. Mutant Kras- and p16-regulated NOX4 activation overcomes metabolic checkpoints in development of pancreatic ductal adenocarcinoma. Nat Commun. Feb 24 2017;8:14437. doi:10.1038/ncomms14437

52. Hall WA, Goodman, KA. Radiation therapy for pancreatic adenocarcinoma, a treatment option that must be considered in the management of a devastating malignancy. Radiation Oncology. 2019/06/26 2019;14(1):114. doi:10.1186/s13014-019-1277-1

53. Liu C-X, et al. Adenanthin targets peroxiredoxin I and II to induce differentiation of leukemic cells. Nature chemical biology. 2012;8(5):486.

54. Jia W, Chen P, Cheng, Y. PRDX4 and its roles in various cancers. Technology in Cancer Research & Treatment. 2019;18:1533033819864313.

55. Mizutani K, et al. The impact of PRDX4 and the EGFR mutation status on cellular proliferation in lung adenocarcinoma. International journal of medical sciences. 2019;16(9):1199.

56. Kim TH, et al. Suppression of peroxiredoxin 4 in glioblastoma cells increases apoptosis and reduces tumor growth. PloS one. 2012;7(8)

57. Medrano M, et al. Interrogation of Functional Cell-Surface Markers Identifies CD151 Dependency in High-Grade Serous Ovarian Cancer. Cell Rep. Mar 7 2017;18(10):2343–2358 doi:10.1016/j.celrep.2017.02.028

58. Johnson WE, Li C, Rabinovic, A. Adjusting batch effects in microarray expression data using empirical Bayes methods. Biostatistics. 2006;8(1):118–127 doi:10.1093/biostatistics/kxj037

59. O'Kane GM, et al. GATA6 Expression Distinguishes Classical and Basal-like Subtypes in Advanced Pancreatic Cancer. Clinical cancer research : an official journal of the American Association for Cancer Research. 2020/03// 2020;doi:10.1158/1078-0432.ccr-19-3724

60. Wiederschain D, et al. Single-vector inducible lentiviral RNAi system for oncology target validation. Cell cycle (Georgetown, Tex). Feb 1 2009;8(3):498–504 doi:10.4161/cc.8.3.7701

61. Therneau TM, Grambsch, PM. The Cox Model. In: Therneau TM, Grambsch PM, eds. Modeling Survival Data: Extending the Cox Model. Springer New York; 2000:39–77.

62. Therneau T. A package for survival analysis in R. 2020;

63. Wieckowski MR, Giorgi C, Lebiedzinska M, Duszynski J, Pinton, P. Isolation of mitochondria-associated membranes and mitochondria from animal tissues and cells. Nat Protoc. 2009;4(11):1582–90 doi:10.1038/nprot.2009.151

